# Alternating Access of a Neurotransmitter:Sodium Symporter Bacterial Homolog Determined from AlphaFold2 Ensembles and DEER Spectroscopy

**DOI:** 10.1101/2024.04.08.587957

**Authors:** Alexandra C. Schwartz, Richard A. Stein, Eva Gil-Iturbe, Matthias Quick, Hassane S. Mchaourab

## Abstract

Neurotransmitter:sodium symporters (NSSs) play critical roles in neural signaling by regulating neurotransmitter uptake into cells powered by sodium electrochemical gradients. Bacterial NSSs orthologs, including MhsT from *Bacillus halodurans*, have emerged as model systems to understands the structural motifs of alternating access in NSSs and the extent of conservation of these motifs across the family. Here, we apply a novel computational/experimental methodology to illuminate the energy landscape of MhsT alternating access. Capitalizing on our recently developed method, Sampling Protein Ensembles and Conformational Heterogeneity with AlphaFold2 (SPEACH_AF), we derived clusters of MhsT models spanning the transition from inward-facing to outward-facing conformations. Systematic application of double electron-electron resonance (DEER) spectroscopy revealed ligand-dependent movement of multiple structural motifs that underpins MhsT’s conformational cycle. Remarkably, comparative DEER analysis in detergent micelles and lipid nanodiscs highlight the profound effect of the environment on the energetics of conformational changes. Through experimentally-derived selection of collective variables, we present a model of ion and substrate powered transport by MhsT consistent with the conformational cycle derived from DEER. Our findings not only advance the understanding of MhsT’s function but also uncover motifs of conformational dynamics conserved within the broader context of the NSS family and within the LeuT-fold class of transporters. Importantly, our methodological blueprint introduces a novel approach that can be applied across a diverse spectrum of transporters to describe their energy landscapes.

**Significance Statement:** The neurotransmitter:sodium symporter (NSS) family plays a crucial role in neurotransmitter reuptake, a sodium-dependent process that transports neurotransmitters from the synapse back into the neuron. This study investigates the bacterial tryptophan transporter MhsT, a homolog of human NSSs, using the deep learning method AlphaFold2 in conjunction with double electron-electron resonance spectroscopy. This combined approach enables us to map the energy landscape that dictates the conformational shifts crucial for MhsT’s function. Furthermore, we reveal how the environment modulates the transporter’s dynamics. From our research, we develop a model of MhsT transport that highlights the extent of mechanistic conservation across the NSS family. Additionally, we introduce a comprehensive framework for exploring the energetic landscapes of transporters, effectively integrating computational and experimental methods.

## Introduction

Neurotransmitter:sodium symporters (NSSs) couple the electrochemical sodium gradient to the thermodynamically uphill transport of neurotransmitters. The NSS family notably includes transporters of neurotransmitters such as serotonin (SERT), dopamine (DAT), and norepinephrine (NET), which are implicated in a variety of neuropsychiatric disorders in humans (1, 2). They are also molecular targets for psychoactive drugs that include multiple classes of antidepressants as well as addictive substances such as cocaine and amphetamines (3–7). Given their medical significance, there has been a continuous effort to understand the underlying functional mechanisms of NSSs, including their bacterial homologs (8–10), with a focus on the alternating access model. This model, central to the complex picture of transport, describes the conformational changes driven by the sequential or coupled binding of ions and substrates (11, 12). Such changes permit the transporter to alternately expose the substrate-binding site to the interior and exterior of the cell, thus enabling substrate uptake against its concentration gradient.

The basic architecture of NSSs was defined in 2005 with the release of the crystal structure of bacterial leucine transporter, LeuT (13), which established it as the archetype model for studying NSSs. The “LeuT fold” is characterized by an inverted structural repeat between TMs 1-5 and TMs 6-10. Interpreted within the context of alternating access, early LeuT crystal structures captured outward-occluded and outward-facing (OF) conformations (13, 14). However, due to its low rate of transport associated with the energetically unfavorable transition from the occluded to inward-facing (IF) conformation (15), the presence of antibodies and/or mutations was required to stabilize LeuT in IF conformations (16, 17). Though earlier studies suggested mechanistic conservation between LeuT and other NSSs (18), more recent cryo-EM structures of hSERT in occluded and IF conformations have revealed differences in their energy landscapes and corresponding mechanisms (19, 20). These distinctions challenge the suitability of LeuT as a model for identifying conserved features and gaining evolutionary insights into the relationship between bacterial and mammalian NSSs.

MhsT is a hydrophobic amino acid transporter from *Bacillus halodurans* that binds and transports tryptophan with an affinity and rate constant comparable to hSERT (3, 21–23), which contrasts LeuT whose transport rate is three orders of magnitude slower than mammalian NSSs (24). Several X-ray crystal structures have been determined for MhsT in the presence of multiple hydrophobic amino acids, all designated as capturing an occluded IF state (21, 25). In this conformation, MhsT typically adopts a partially unwound intracellular TM5 (TM5i) similar to that observed in hSERT. While the MhsT crystal structures share some similar features with IF hSERT cryo-EM structures, the biological significance of this conformation in the transport cycle has not been established. The assignment of MhsT’s crystal structures as IF initially relied on the presence of a partially unwound TM5i that allows water to access the intracellular vestibule (21). In these structures, the unwound region spanning five residues has an N-terminal helical cap, a structural detail that is absent in hSERT where the unwinding instead elongates the TM4-TM5 loop (21). Interestingly, the lipidic cubic phase (LCP) crystallization method yielded a Na^+^/Trp-bound structure of MhsT that featured a continuously helical TM5 (PDB: 4US4) (21). While the structure is also designated as IF, it deviates from other crystal structures beyond TM5. The N-terminus and TM1a residues at the N-terminal cap are unresolved, indicative of a highly dynamic region. Scaffold TMs 8 and 9, along with bundle TMs 2, 6, and 7, also show shifts away from their respective domains. Computational analysis predicted that both 4US3 and 4US4 structures are sampled as intermediates in the MhsT conformational cycle (26). Reflecting its robust transport rate and homology to mammalian NSSs, MhsT has emerged as a model of choice for applying a broad spectrum of structural biology, computational, and spectroscopic methods to illuminate conserved aspects of transport among NSSs (21, 25–28).

While high-resolution structures provide crucial information for the molecular architecture and ligand coordination, they represent static structures that are extrapolated to generate hypothetical transport mechanisms. In addition to issues with contacts in the lattice, crystal structures capture the lowest energy state under the given set of conditions, which may or may not be frequently sampled during the transport mechanism. Though cryo-EM structures do not have the same level of conformational selection as crystal structures, there are some biases associated with sample preparation, particle picking, and model construction. Universal to all structural and biophysical techniques, the purification conditions—especially the choice of detergents or lipids used—can bias protein conformations and alter their energetics (27, 29–31), a crucial aspect that will be further explored in this paper.

In this study, we introduce a novel methodological approach to investigate the structural dynamics associated with MhsT ligand binding and its lipid dependence by combining Double Electron-Electron Resonance (DEER) spectroscopy with an Alphafold2 (AF2) (32, 33) method, Sampling Protein Ensembles and Conformational Heterogeneity (SPEACH_AF) (34). Our MhsT DEER data were evaluated in both n-decyl-β-maltoside (DM) detergent micelles and lipid nanodiscs. We found that these two environments illuminate distinct aspects of MhsT’s conformational dynamics. Ligand-dependent DEER distance distributions and AF2 conformational ensembles were integrated to outline the energetic landscape of MhsT alternating access. Comparisons to hSERT’s cryo-EM structures, alongside structural motifs in LeuT and the LeuT-fold transporter Mhp1 assessed by crystal structures and DEER, reveal conserved and divergent elements of alternating access. Furthermore, the results emphasize the distinct energetics of conformational states as factors underlying the mechanistic diversity between these transporters.

## Results

### Methodology

To map ligand-dependent alternating access in MhsT, spin label selection was guided by hSERT cryo-EM structures and our previous DEER studies of LeuT and Mhp1 to target TMs proposed to mediate conformational changes (19, 20, 31, 35). NSS nomenclature designates TMs 1, 2, 6, and 7 as the bundle domain and TMs 3, 4, 8, and 9 as the scaffold domain (9, 16). In multiple LeuT-fold transporters, TMs 5 and 10 are implicated in the transport mechanism and are sometimes referred to as the gating helices (32). Extracellular spin label pairs included two sites on the scaffold (TMs 4 and 8) to be used as references, sites on TMs 1b, 6a, and 7 to evaluate the movement of the bundle, and sites on TMs 5 and 10 to evaluate their role as gating TMs (Fig. 1A). Intracellular spin label sites were originally chosen to interrogate movement of the bundle (TMs 1a, 2, and 6b) and gates (TMs 5 and 10) using a scaffold label site (TM9) as a reference (Fig. 1B). However, due to limited accessibility of intracellular TM6b and TM10, their movements were estimated by corroborating spin labels on adjacent intracellular TM7 and TM11, respectively (Fig. 1B). DEER data for spin label “pyramids”, which consist of all six combinations of four sites, can be used to extract information regarding each site’s location and movement (Fig. 1). The extracellular and intracellular spin label pyramids consist of TMs 1b, 4, 5, and 8 (Fig. 2 *A* and *B*) and TMs 1a, 2, 9, and 11 (Fig. 3 *A* and *B*), respectively. All spin labeled MhsT mutants were verified for transport activity by measuring radiolabeled substrate uptake into proteoliposomes (*SI Appendix,* Fig. S1 and Table S1).

**Fig. 1.**
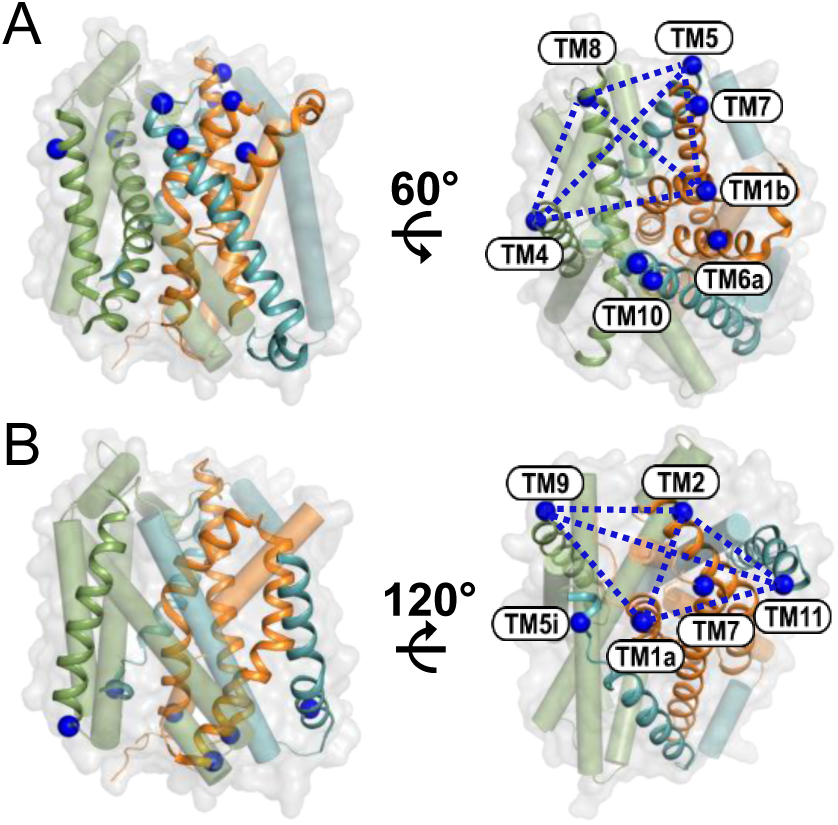
MhsT spin label sites. Cartoon of Na+/Trp-bound crystal structure of MhsT (PDB: 4US3) with label sites depicted as blue spheres. The scaffold domain (TMs 3, 4, 8, 9) is colored green while the bundle domain (TMs 1, 2, 6, 7) is shown in orange. Gating helices TMs 5 and 10 as well as TM11 are shown in teal. Extracellular (A) and intracellular (B) label sites on TM helices, shown from the side (Left) and from their respective sides with the spin label pyramids outlined by dashed blue lines (Right). TMs without label sites are shown as cylinders.

**Fig. 2.**
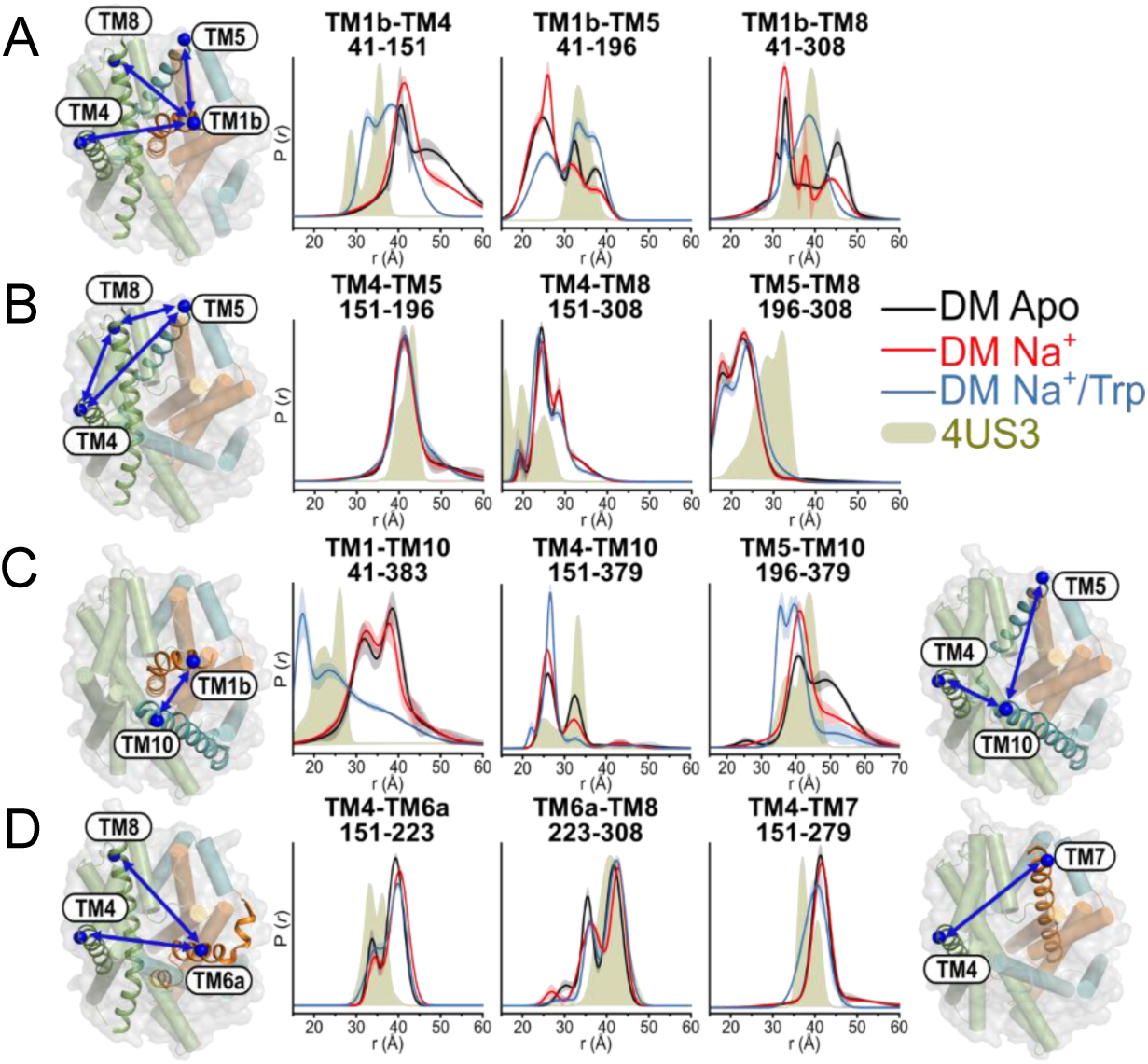
Closure of the extracellular side in detergent micelles is characterized by ligand-dependent movements of TM1b and TM10. Cartoon depictions as shown in Fig. 1A, featuring blue arrows to represent distances between spin label sites. Distance distributions are shown under apo (black), Na+ (red), and Na+/Trp (blue) conditions, with their respective confidence bands. Predicted distance distributions from MhsT crystal structure (PDB: 4US3) are shown as solid gold. The extracellular pyramid reveals ligand-dependent changes in TM1b (A) while TMs 4, 5, and 8 are remain relatively static (B). TM10 acts as a gate with TM1b in the extracellular closure mechanism (C). TM6a does not undergo changes between conditions (D, Left) while Na+/Trp induces a minor rearrangement of TM7 (D, Right).

**Fig. 3.**
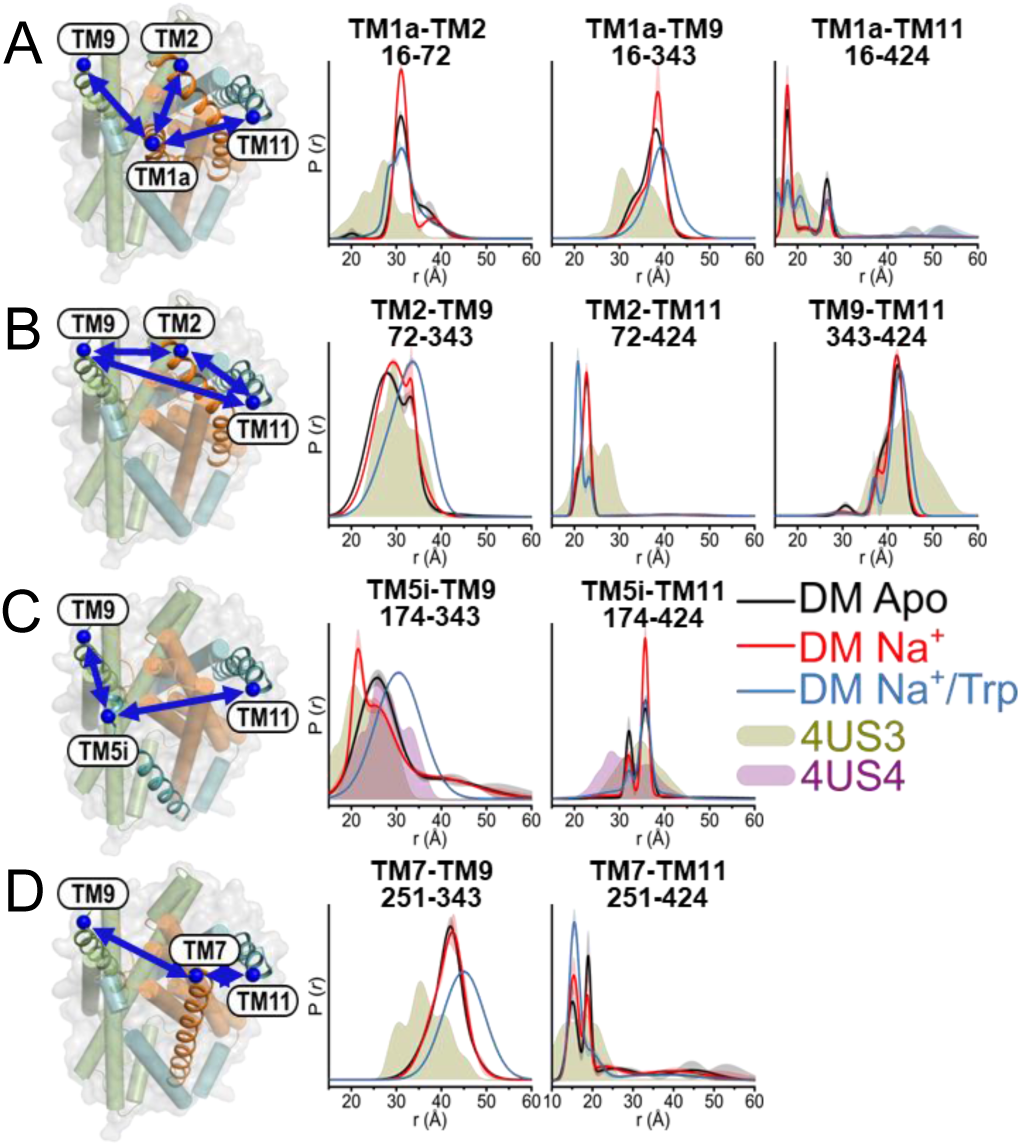
MhsT’s intracellular opening in detergent micelles is defined by movements of the bundle and TM5i. Cartoon depictions of label sites are as shown in Fig. 1B. Experimentally determined and predicted distributions are as depicted in Fig. 2. The intracellular pyramid reveals that TM1a undergoes ligand-dependent changes between conditions (A) while TM2 shifts in the presence of Na+/Trp (B, Left) as TMs 9 and 11 remain relatively static (B, Right). TM5i acts as a gate with TM1a (C), shown with predicted distributions for the fully helical TM5 structure (solid purple). TM7 also undergoes Na+/Trp-induced changes (D).

Previous studies showed that MhsT displays similar binding activity in DM detergent as in nanodiscs (27), which was interpreted to suggest that MhsT conformational dynamics would be similar in both environments. Therefore, the initial set of experiments was performed in DM detergent (*SI Appendix*, Figs. S2 and S3), and conformational states were assigned based on analysis of the distance components observed in DEER data. Pairs of interest were reconstituted into nanodiscs and evaluated for differences in ligand-dependent conformational dynamics (*SI Appendix*, Figs. S4 and S5). For both environments, three biochemical conditions were examined: in absence of ions and substrate (apo), Na^+^, and Na^+^/Trp. The continuous-wave electron paramagnetic resonance (CW-EPR) spectra of the spin-labeled mutants (*SI Appendix*, Figs. 2-5, *Right*) under each biochemical condition did not reveal variations in mobility, suggesting that ligand-dependent distance changes are not attributed to rotameric changes. Therefore, we interpret the changes in distance components as reflecting shifts between conformations. Computational models were generated using SPEACH_AF and compared to experimental data. Predicted spin-label pair distributions for crystal structures and computational models were simulated with ChiLife using MMM methodology (37, 38).

**Fig. 4.**
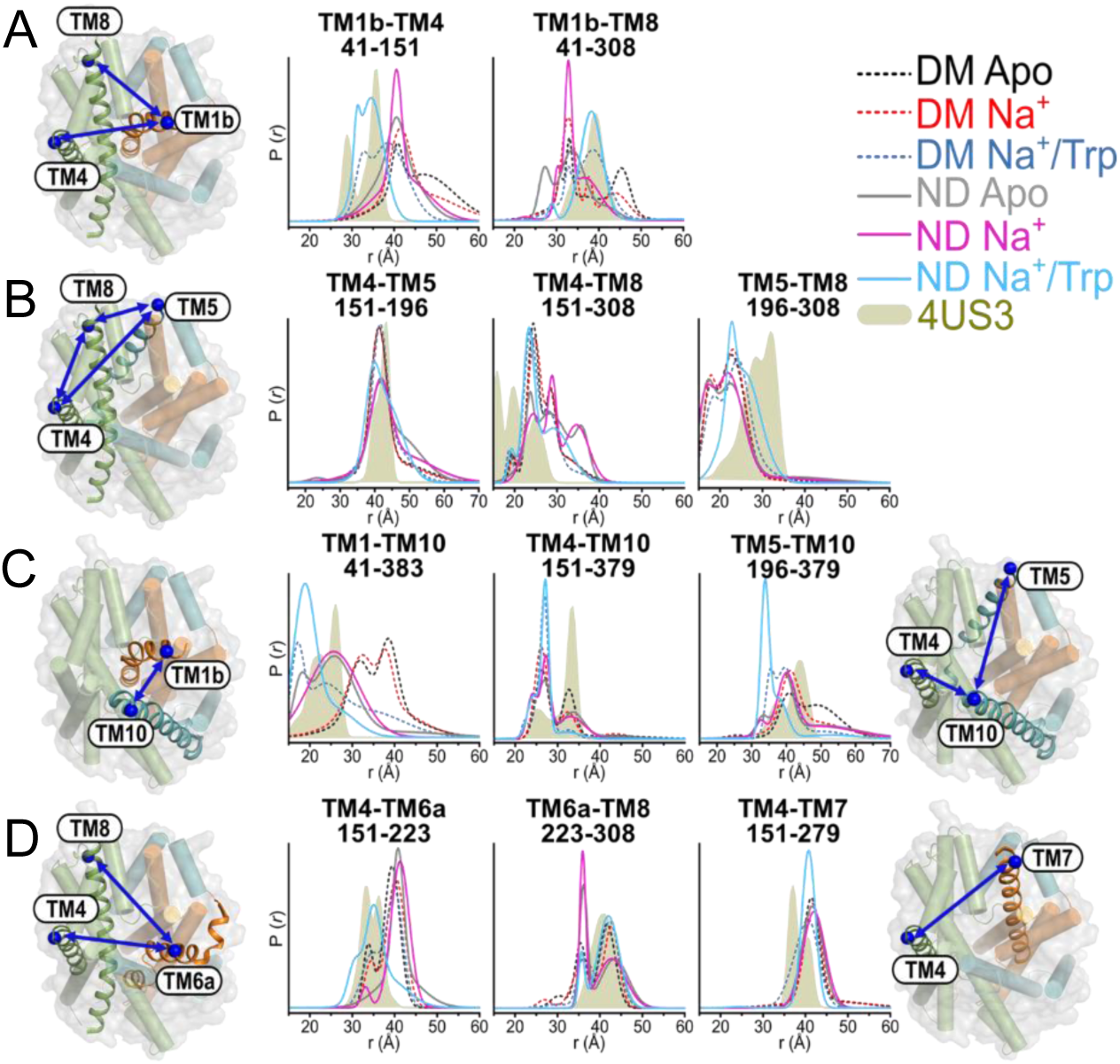
The extracellular side of MhsT undergoes ligand-dependent changes in lipid nanodiscs not observed in detergent micelles. Spin label pairs are depicted as in Fig. 2. Distance distributions for pairs in nanodiscs (solid lines) are shown under apo (gray), Na+ (pink), and Na+/Trp (light blue) conditions. Detergent distributions (dashed lines) are colored as in Fig. 2. Predicted distance distributions from the crystal structure (PDB: 4US3) are shown as solid gold. TM1b (A) and TM10 (C) exhibit more pronounced Na+/Trp-induced effects in nanodiscs compared to detergent micelles. In nanodiscs, the presence of Na+/Trp induces changes in TM5 and TM8 distributions (B), shifts TM6a towards the scaffold (D, Left), and has a reduced effect on TM7 relative to detergent micelles (D, Right).

**Fig. 5.**
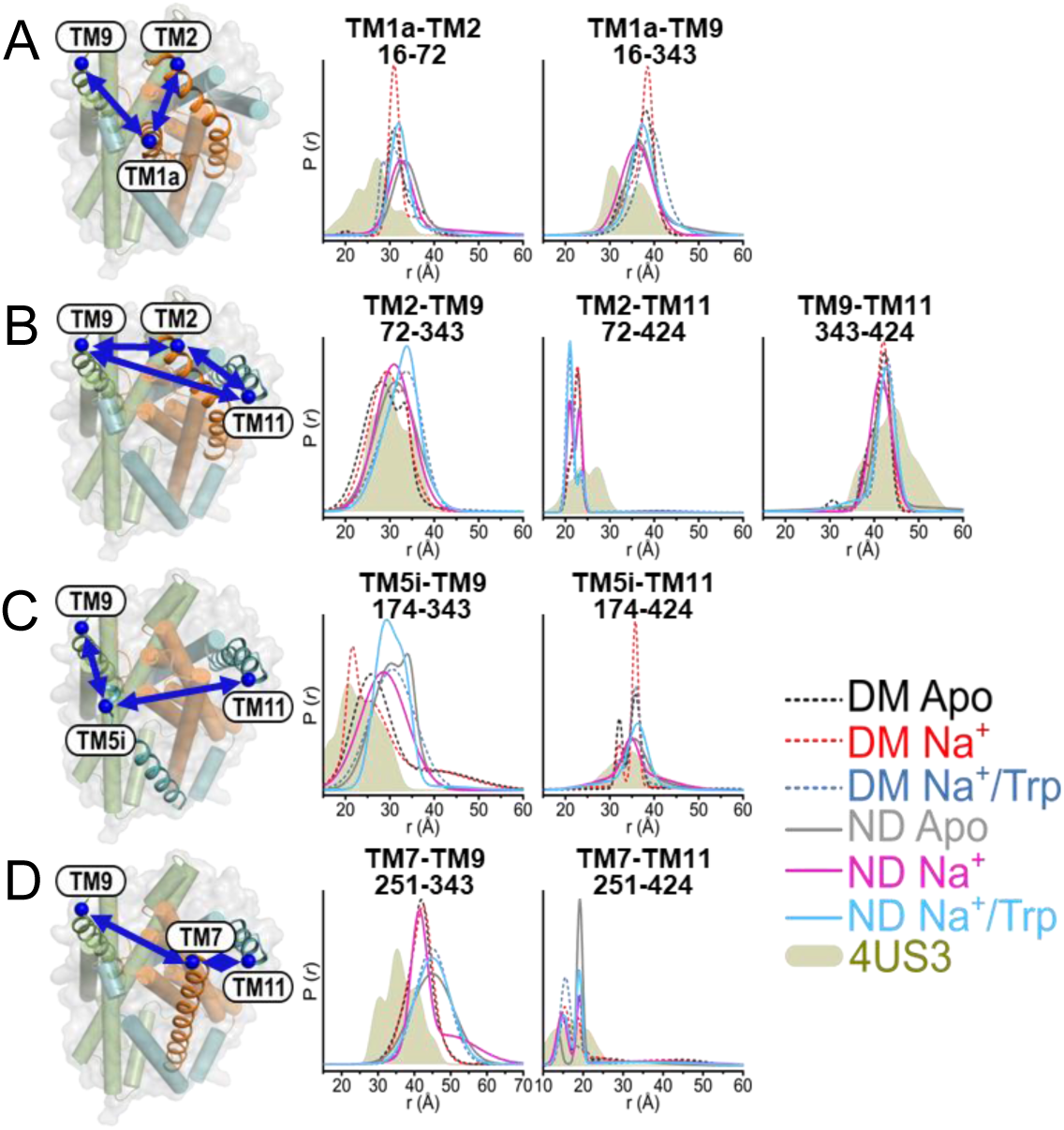
Lipid bilayers enhance the dynamics of the intracellular side of MhsT compared to detergent micelles. Spin label pairs are depicted as in Fig. 3. Experimentally determined and predicted distributions are as indicated in Fig. 4. The magnitude of ligand-dependent effects for TM1a (A) and TM5i (C) are reduced in nanodiscs.

### Closure of the extracellular side in detergent micelles entails coordinated movement of TM1b and TM10

TM1b is involved in coordinating the ion and substrate binding sites as well as the extracellular gate, features that are conserved across NSSs (39). The DEER data, deduced from the extracellular pyramid that includes TMs 1b, 4, 5, and 8, paints a picture of TM1b as a highly dynamic helix (Fig. 2 *A* and *B*). Multicomponent distance distributions suggest TM1b adopts several conformations sampled under equilibrium between the three biochemical conditions. The absence of ions and substrate, i.e. apo condition, promotes enhanced conformational sampling that deviates from the IF crystal structure (4US3, Fig. 2, solid gold), whereas the addition of Na^+^ stabilizes more unique states as evidenced by the narrowed distance distribution. In particular, Na^+^ binding appears to steer TM1b toward the scaffold domain. Notably, Trp induces large structural rearrangements of TM1b that align with the outward-occluded conformation identified by the crystal structure.

While both TM1b and TM10 are involved in forming a conserved salt bridge that acts as an extracellular gate for NSSs (39), the role of TM10 in the alternating access mechanism is not conserved to the same extent as TM1b (36). With reference to TMs 4 and 5, TM10 samples multiple conformations in the apo condition (Fig. 2C). Similar to TM1b, the presence of Na^+^ and Trp promotes conformational selection and movement of TM10 toward the bundle domain. Interestingly, the movement of TM10 is not directly coupled to TM1b as there is a large change in the distance distribution for the TM1b to TM10 distance in the presence of Na^+^/Trp (Fig. 2C). While the crystal structure predictions match the Na^+^/Trp distributions for TM1b pairs, it only partially agrees with the TM10 Na^+^/Trp distributions.

Surprisingly and in contrast to TM1b and TM10, other structural changes between the bundle and scaffold domains are largely muted or absent in detergent micelles. Significantly, TM6a does not exhibit ligand-dependent changes (Fig. 2D). A minor rearrangement of TM7 induced by Trp binding was observed. We note that TM7 has conserved roles in Na1 binding and as an intracellular gate in mammalian NSSs (39), but residues mediating these roles are not conserved in MhsT. Importantly, lack of distance changes in TM8 (Fig. 2B) suggests that substrate-dependent conformational changes in TM7 are not propagated through EL4 as determined for other LeuT-fold transporters (31, 35, 40, 41).

Overall, the extracellular data in detergent micelles suggests the presence of three conformations characterized by movements of TMs 1b and 10 towards the scaffold, suggesting extracellular closure in the presence of Na^+^/Trp. Coordinated movement of TM1b and TM10 is emphasized by the largest amplitude change in distance (up to 20 Å) between the helices (Fig 2C).

### TM1a and TM5i underpin opening of the intracellular side in detergent micelles

In detergent micelles, TM1 acts alongside gating TM5 to regulate the transporter’s access to both sides. On the intracellular side, TM1a and TM5i undergo the largest ligand-dependent changes in MhsT as evidenced by the DEER distance distributions, demonstrating their importance in the inward opening mechanism. TM1a’s movement relative to other sites in the intracellular spin label pyramid, TMs 2, 9, and 11 (Fig. 3 *A* and *B*), reinforces its conserved role in the intracellular opening across NSSs (34, 37). Multi-component distributions characterize TM1a conformational dynamics as manifested in all three spin label pairs (Fig. 3A). When bound to Na^+^/Trp, a movement of TM1a away from the scaffold is evident from the TM1a-TM9 distribution, signifying intracellular opening. The appearance of additional distance components implies that TM1a dynamics increase with Trp binding. Despite their width being overestimated (38), the predicted distributions from the crystal structure do not closely correspond with one particular component across the TM1a pairs, indicating a deviation between TM1a’s position in the crystal structure and in solution.

Whereas TM5’s role in the intracellular opening mechanism has been recently highlighted in MhsT and hSERT (19, 21, 26), its conservation across LeuT-fold transporters has not been experimentally evaluated thoroughly (36, 39). The TM5i label site was intentionally placed on a helical segment to avoid changes in spin label rotamers associated with the putative backbone unwinding, which could convolute interpretation of the DEER data. CW-EPR spectra of the singly labeled TM5i site (174C) confirm its placement in an ordered region of TM5i (*SI Appendix,* Fig. S6), indicating the spin label is not excessively mobile and has similar mobility between the three conditions. Nevertheless, multiple distance components in TM5i pairs are observed, suggesting multiple conformations of TM5i (Fig. 3C). Similar to TM1a, the addition of Trp promotes large-scale movement of TM5i away from the scaffold while concomitantly increasing TM5i dynamics as evidenced by increased breadth of the distribution. Comparison of the predicted distances from the Na^+^/Trp-bound crystal structures with the fully wound TM5i (4US4) and partially unwound TM5i (4US3) reveals a similar pattern of TM5i movement captured by the apo and Na^+^ distributions (Fig. 3C). While this trend could support TM5i unwinding under the Na^+^ condition in detergent micelles, it is possible that other TM5i movements could result in similar shifts in distribution.

In addition to the primary ligand-dependent changes observed in TMs 1a and 5i, TMs 2 and 7 undergo movements in the presence of Na^+^/Trp (Fig. 3 *B* and *D*). More pronounced with reference to TM9, both TM2 and TM7 shift away from the bundle domain in a comparable fashion to TM1a and 5i. Moreover, the broadened distributions suggest increased dynamics of TM2 and TM7, in keeping with the pattern observed with TM1a and TM5i. Collectively, these data emphasize the coupled movement of helices in the bundle domain away from the scaffold to establish an IF intermediate driven by Trp binding.

### Lipid bilayers consolidate changes between apo and Na^+^ conditions and couple extracellular TM movements

Despite previous biochemical data suggesting similar behavior of MhsT in DM micelles and lipid bilayers (27), reconstitution into nanodiscs (3:1 *Escherichia coli* polar lipids: egg phosphatidylcholine) exposes dramatic differences in MhsT’s conformational dynamics. In general, lipids reduce large amplitude dynamics in the extracellular gating TMs yet enhance the dynamics of other TMs, exposing coupled movements on the extracellular side.

Detailed inspection of the DEER data indicates that although TM1b remains a major element in the extracellular closure mechanism, the populations of sampled components for each condition change substantially relative to detergent (Fig. 4A). For example, the lipid environment suppresses the population of the longest-distance components of TM1b-TM4 and TM1b-TM8 pairs in the absence or presence of Na^+^. As a result, the TM1b distributions under the apo and Na^+^ conditions are more comparable in nanodiscs than in detergent. Na^+^/Trp shifts TM1b towards the same populations as in detergent but with a more pronounced effect in nanodiscs.

TM5 (gating) and TM8 (scaffold) undergo ligand-dependent changes in lipid nanodiscs that are not observed in detergent micelles (Fig. 4B). Relative to TM4, extracellular TMs 5 and 8 sample longer distances under the apo and Na^+^ conditions to a greater extent than observed in detergent. This suggests that TM5 and TM8’s dynamics are restricted in detergent micelles. Moreover, the substantial distance increase in the TM5-TM8 distribution in the presence of Na^+^/Trp further supports the relative movement of TM5.

While TM10 undergoes ligand-dependent changes between all three biochemical conditions in detergent micelles, the effect of Na^+^/Trp is accentuated in nanodiscs as the changes between the apo and Na^+^ distributions are abrogated (Fig. 4C). However, the long-distance components associated with the apo distributions in detergent are not substantially reduced as observed with TM1b but rather broaden in nanodiscs, suggesting a more dynamic TM10 conformation compared to detergent micelles. Stabilization of shorter distance components between TM4-TM10 and TM5-TM10 relative to detergent demonstrates that the effect of Na^+^/Trp on TM10 is amplified in nanodiscs, in a similar manner to TM1b.

In contrast to detergent micelles, TM6a appears to participate in a collective bundle movement in nanodiscs that is driven by Trp binding (Fig. 4D, Left). This is evident in the TM4-TM6a pair where conformational changes are mitigated by detergent. Although only a relatively minor distance component in the detergent distributions, Trp binding induces a nearly complete ∼5Å shift towards the scaffold as observed with TM1b. An additional underlying broad distance component is present under all three conditions in nanodiscs, suggesting that TM6a is more dynamic and experiences a higher range of fluctuations compared to detergent micelles. Notably, the predicted crystal structure distributions align with the Na^+^/Trp distributions for both TM6a pairs in nanodiscs. This observation implies that TM6a is involved in MhsT’s extracellular closure mechanism, which agrees with conserved TM6a movements observed in other NSS structures (18, 28).

Whereas Na^+^/Trp broadens the TM4-TM7 distribution in detergent, the TM4-TM7 distribution becomes narrower in nanodiscs (Fig. 4D, *Right*). In both environments, Na^+^/Trp distributions are shifted to shorter distances relative to the apo and Na^+^ conditions, but this effect is reduced in nanodiscs. The reduced conformational sampling for extracellular TM7 and TM8 suggests their movements may be coordinated in nanodiscs, which contrasts observations in detergent micelles. This Na^+^/Trp-induced coordinated movement between TM7 and TM8 likely involves EL4 in nanodiscs.

### Lipid bilayers temper intracellular ligand-dependent distance changes while enhancing dynamics

Whereas reconstitution of MhsT into nanodiscs enforces coordination of the bundle and TM10 movement on the extracellular side, a strikingly distinct effect is observed on the intracellular side (Fig. 5). The ligand-dependent shifts appear dampened relative to the detergent data. Instead, broader distance distributions indicate a more dynamic intracellular side, accompanying shifts in equilibria towards distances characteristic of Na^+^/Trp distributions in detergent. The systemically observed broadened distributions are likely the consequence of enhanced fluctuations between conformations, suggesting that helices comprising the intracellular region may sample a conformational equilibrium with lower energy barriers between intermediates in lipid nanodiscs.

While TM1a undergoes large-scale shifts in detergent micelles, the magnitude of TM1a’s ligand-dependent shifts is significantly reduced in nanodiscs (Fig. 5A). Nanodisc-reconstituted TM1a pairs reveal broad distributions between all three conditions compared to their detergent counterparts. However, the breadth of the TM1a distributions in nanodiscs is ligand-dependent, indicating TM1a is most dynamic under the apo condition and least dynamic in the presence of Na^+^/Trp.

TM2 follows a similar trend of ligand-dependent changes in nanodiscs as observed in detergent, although TM2 shifts to longer distances that are characteristic of an IF conformation under apo and Na^+^ conditions (Fig. 5B). The narrowed Na^+^/Trp distribution in nanodiscs indicates that TM2’s shift towards a more open state may complement extracellular conformational changes observed in nanodiscs.

TM5i undergoes a similar effect to TM2 in the presence of Na^+^/Trp, stabilized further towards the same component observed in detergent micelles (Fig. 5C). Notably, the apo distribution more closely resembles the Na^+^/Trp distribution than the Na^+^ distribution, though its increased breadth suggests enhanced dynamics. Interestingly, the unique movement observed in the presence of Na^+^ in detergent micelles is absent. The width of the Na^+^ distribution extends through the full range of TM5i distances, suggesting that TM5i could adapt any position across the distance range sampled in detergent micelles and lipid nanodiscs. However, the considerable overlap of the Na^+^ distribution with the other distributions in nanodiscs indicates that the conformations stabilized by Na^+^ in detergent are lower probability.

Consistent with the effects observed in TMs 2 and 5i, Na^+^/Trp similarly affects TM7 in both detergent and nanodisc environments. Mirroring TM5i-TM9, the TM7-TM9 distribution under apo and Na^+^ conditions in nanodiscs sample longer distances than in detergent, with the apo distribution more closely resembling that observed with Na^+^/Trp. Overall, these data highlight the enhanced dynamics of MhsT’s intracellular side in nanodiscs, particularly under the apo condition. Although Na^+^/Trp-induced shifts remain evident in nanodiscs, the reduced extent of ligand-dependent changes implies a more significant role of the extracellular side in regulating transport.

### Modeling of MhsT conformational dynamics with SPEACH_AF

To contextualize the DEER data within a conformational landscape of MhsT, we applied SPEACH_AF, an *in-silico* mutagenesis approach that harnesses AF2 to model protein ensembles (34). Briefly, alanine substitutions were introduced across the multiple sequence alignment (MSA) using a sliding window approach. For each mutated segment, the five AF2 modelers generated three models each, resulting in a total of 1,920 models. Models were subsequently filtered by principal component analysis (PCA) and the average pLDDT (34, 42), though all MhsT models passed these filtering steps. Fig. 6A shows the two principal components of the PCA that include the crystal structures. Multiple helical segments undergo relative displacement between these models (see below). Notably, PC1 captures the distinct variation in TM5 where the crystal structures featuring a partially unwound TM5 (PDBs: 4US3, 6YU2, 6UY3, 6YU4, 6YU5, 6YU6, and 6YU7) diverge from 4US4, which possesses a fully helical TM5.

**Fig. 6.**
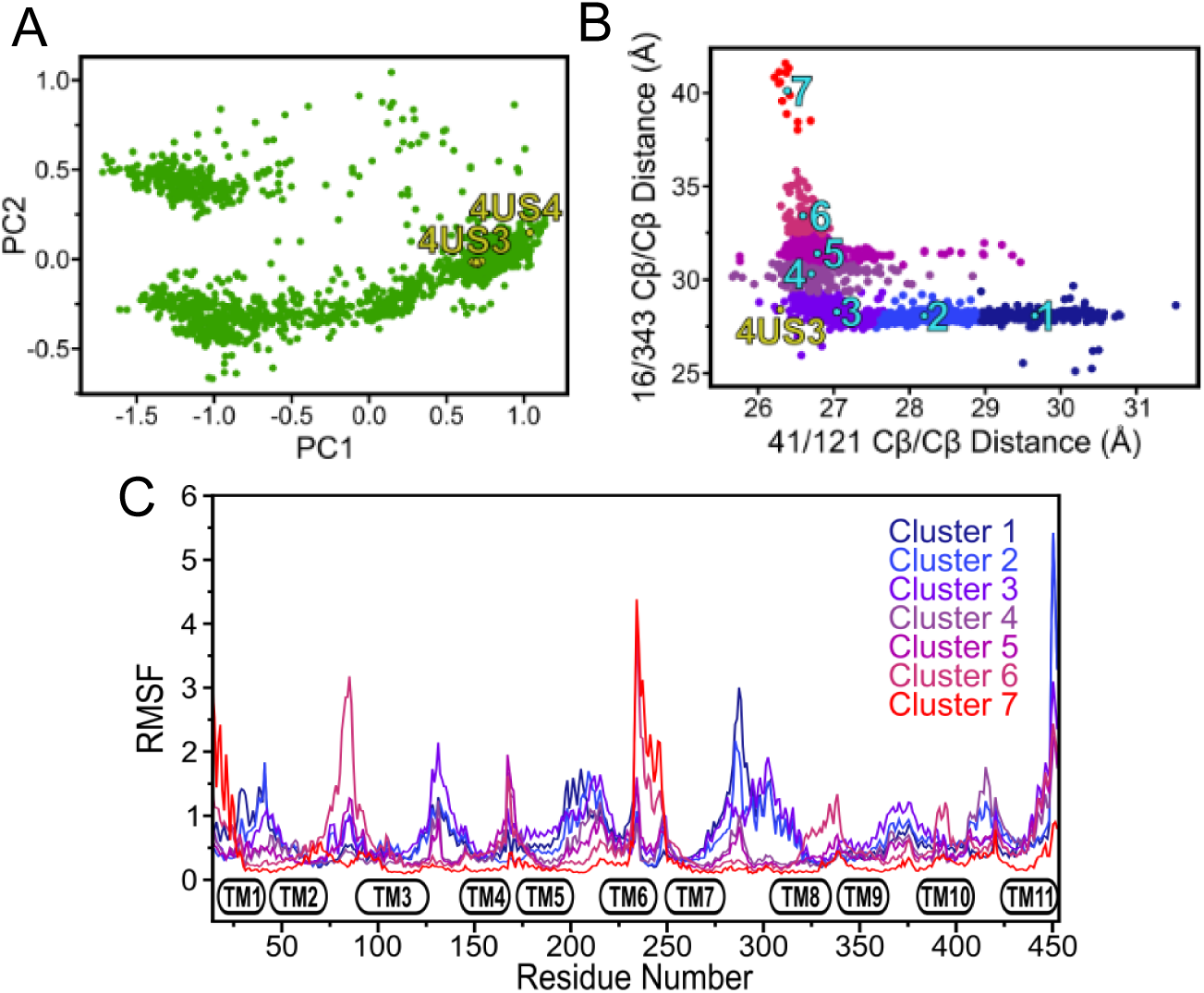
Analysis of models by principal components, collective variable clustering, and intra-cluster variability. (A) The first two principal components for the models (green) generated by SPEACH_AF and the crystal structures (gold). The 4US4 structure with the fully wound TM5 is to the far right compared to the other crystal structures that are marked by a cluster near the 4US3 structure. (B) Collective variable analysis and K-means clustering of the SPEACH_AF models places 4US3 near the interface between the occluded Clusters 3 and 4. (C) Intra-cluster residue fluctuation profiles quantify the variability within each cluster.

To relate the conformational ensembles along PC1 to the experimental distance distributions, we simulated spin-label distances for all of the experimental pairs and plotted the average distance vs. PC1 (*SI Appendix*, Fig. S7). As observed in our benchmark set (34), PC1 captures conformational changes underpinning alternating access. For MhsT, we find that models at low and high PC1 values correspond to OF and IF conformations, respectively. The observed agreement in the trends between simulated and experimental distances suggests that the AF2 conformational ensemble reflects the direction of ligand-dependent changes, though the values are not directly comparable due to intrinsic uncertainties in spin-label modeling (38, 43). Interestingly, the changes on the extracellular side span the whole PC1 space, whereas changes on the intracellular side are confined to a smaller range of PC1 space. This suggests an asymmetry in the extent of conformational changes across the two sides of the transporter with more disorder on the extracellular side.

We therefore explored collective variables to more directly capture structural changes on both sides of the transporter. First, the per-residue RMSF was calculated across all models, which emphasized that the extracellular side is predicted to have greater mobility compared to the intracellular side (*SI Appendix,* Fig. S8). Next, we characterized regions by high and low mobility and examined pairs of residues exhibiting changes reflective of the entire conformational landscape. The models suggest TM1’s role as a gate across both sides of the transporter, aligning with experimental observations captured by DEER (Figs. 2*A* and 3*A*). Consequently, we monitored the degree of closure of the intracellular and extracellular sides for all the models using pairs 16-343 and 41-121, respectively (Fig. 6B). Based on the generated models and pairs used to parameterize intracellular and extracellular closure, the crystal structures appear to correspond to doubly occluded conformations with both sides closed.

Variations of the two collective variables across models were then clustered in seven groups by K-means analysis (Fig. 6B). To examine the overall conformational space of each cluster, the intra-cluster RMSF was measured (Fig. 6C). Cluster 1 and Cluster 7 represent fully OF and fully IF conformations, respectively. The remaining clusters capture intermediate states that transition from OF to IF conformations. Structural heterogeneity, illustrated in putty representations identify disordered regions primarily found in TM1b, TM5, TM6a, and intracellular loops (*SI Appendix*, Fig S9).

Representative structures from each cluster were selected on the basis of their similarity to other models within the cluster (*SI Appendix*, Fig. S10). Direct comparisons of the representative models, on the basis of per-residue RMSD (Fig. 7), identify hypothetical “hot spots” for conformational changes between clusters. Remarkably, the transition from Cluster 6 to Cluster 7 reveals the most dramatic movements observed across all cluster transitions, primarily occurring in TM1a and TM6b.

**Fig. 7.**
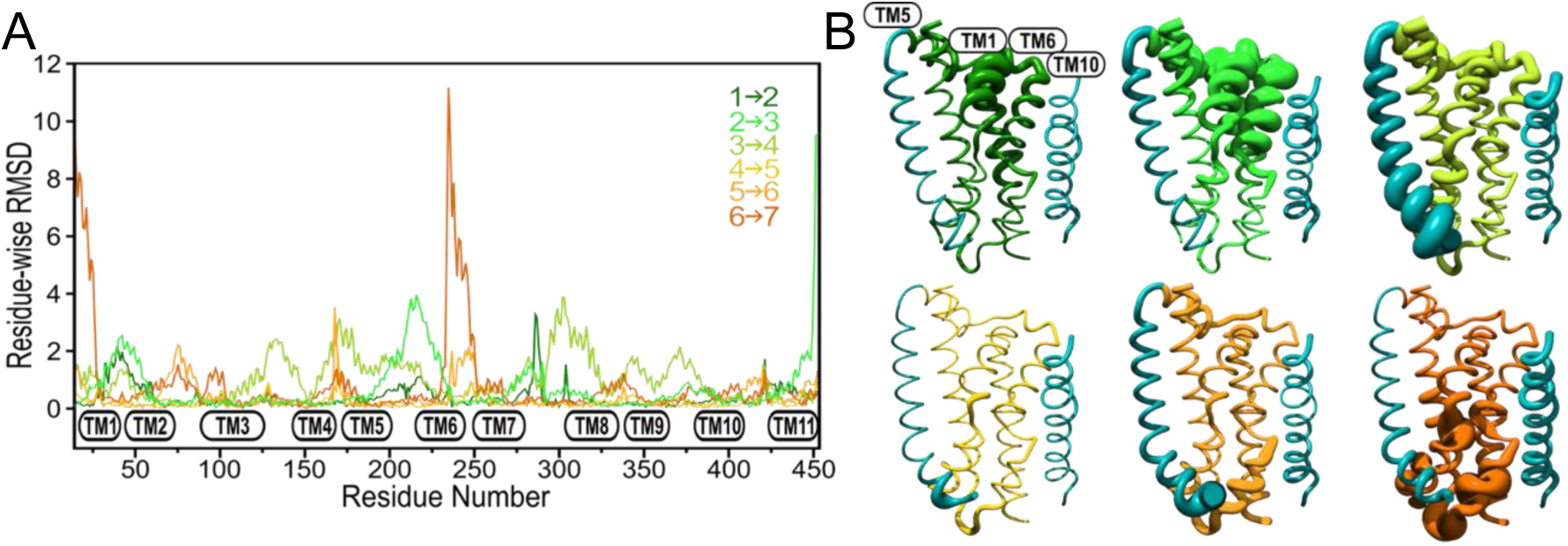
RMSD analysis of implied conformational dynamics of SPEACH_AF clusters from the OF to IF transition. (A) Per-residue RMSD values are plotted to indicate movements of TMs and loop regions as the clusters progress from the most OF (Cluster 1, dark green) to the most IF (Cluster 7, dark orange). (B) RMSD values are mapped onto the initial structural model for each cluster pair transition, emphasizing regions of significant conformational changes and visually tracing the structural evolution from dark green (OF) to dark orange (IF). Gating helices TMs 5 & 10 are highlighted in teal while the scaffold domain along with TM11 are omitted for clarity.

### Alternating access of MhsT, deduced from a comparison of SPEACH_AF clusters with DEER distance distributions

Due to the limited resolution of the DEER data, the experimentally sampled conformations cannot be directly matched to individual structural models generated by SPEACH_AF. Consequently, the model clusters were categorized based on their conformational states: Clusters 1 and 2 were categorized as OF, Clusters 3-5 as occluded, and Clusters 6 and 7 as IF. To better represent the transporter dynamics, the predicted distance distributions for each category were averaged, producing composite distance distributions. Because the DM apo and Na^+^/Trp distributions encompass the extensive range of experimentally observed movements, they were compared to the predicted composite distance distributions (*SI Appendix*, Fig. S11). On the extracellular side, OF models sample longer distances in the TM1b and TM10 pairs, implying a coordinated movement towards the scaffold as the gates close. This pattern is reflected in the DEER data, confirming the transition from an OF to an occluded conformation. While TM6a does not undergo movements associated with extracellular closure in the DM data, its coupled movement towards the scaffold with TM1b in nanodiscs is consistent with the closure suggested by the predicted distributions.

Similarly, on the intracellular side, TM1a and TM5i are closer to the scaffold in the OF conformation, reflected in the shorter distances than those observed in the IF conformation. This variation in distance is essential for enabling the opening necessary for ion and substrate release. TM2 and TM7, according to the SPEACH_AF-predicted distributions, also show distinct distance changes between conformations. These combined observations from the computational and experimental data highlight the intricate conformational dynamics within the bundle domain, validating the predicted conformational shifts by SPEACH_AF and the dynamic landscape captured by DEER measurements.

## Discussion

In this study, we investigated the conformational dynamics of the LeuT-fold, bacterial NSS homolog, MhsT using a combination of DEER experiments and an AF2 methodology developed in our laboratory, SPEACH-AF. Central to MhsT’s mechanism, our data highlight how the lipid bilayer shapes and modulates the energetics of coupled ligand-dependent movements. In the absence of high-resolution conformational intermediates, we introduced a cluster analysis derived from collective variables to parse and select predictive AF2 models that fit with the spectroscopic data. This integrative approach stimulates a structure-and dynamics-based mechanism of MhsT transport and provides a perspective into the conservation of conformational dynamics within the NSS family and across LeuT-fold transporters.

### Lipid bilayers are required to reveal the coupled structural motifs underpinning MhsT conformational dynamics

Although detergent micelles are thought to be more conducive to protein dynamics due to their less rigid structure relative to lipid bilayers, the scale of the TM movements observed in MhsT in detergent micelles was somewhat unexpected. We surmise that these movements expose higher energy intermediates that may be critical for the transport mechanism, despite being less frequently sampled. For instance, the large movement of TM1b that establishes an OF state is less pronounced in nanodiscs and is captured within SPEACH_AF cluster 1 (*SI Appendix*, Fig. S10). Similarly, the IF intermediate, formed in part by large-scale movements of TM1a, is also strongly reduced in nanodiscs. Importantly, cluster analysis of AF2 models captures these changes in TM1a. Generally, we observed that while Na^+^/Trp induces a more distinct shift in extracellular conformational equilibria in nanodiscs, the coupled distance changes on the intracellular side are more subtle and less resolved in otherwise broad distributions that reflect enhanced conformational sampling.

Critical to elucidating conformational dynamics, we discovered that the native-like environment of lipid bilayers is required to identify coupled structural motifs. The structural motifs associated with extracellular closure are amplified as TM5, TM6a, and TM8 undergo ligand-dependent changes in nanodiscs that are minimal or absent in detergent micelles. TM8 displays enhanced dynamics in nanodiscs under apo and Na^+^ conditions that are reduced in the presence of Na^+^/Trp. Closure of the extracellular side was marked by a coordinated movement between TM1b and TM6a as observed in other NSSs (16, 19, 31). Molecular dynamics simulations have suggested that TM1b and TM6a may undergo independent movements in hSERT (44). When bound to Na^+^ and serotonin *in silico*, hSERT can proceed towards a partially occluded intermediate where TM1b completely transitions to the closed position while TM6a fluctuates between the open and partially occluded conformations (44). While this effect is only observed in detergent micelles, our DEER data in detergent micelles experimentally, for the first time, that TM1b and TM6a can move in a non-coordinated fashion in NSSs.

### TM5i dynamics in MhsT and NSSs

The structural record posits that intracellular opening and Na^+^ release are facilitated by the gating dynamics of TM5i during the OF to IF transition (26). In DM micelles, we found that TM5i acts as an intracellular gate, undergoing distinct changes that depend on ligand binding. In particular, the short TM5i-TM9 distance sampled under the Na^+^ condition demonstrates the most agreement with the predicted distance arising from a partially unwound helix (PDB: 4US3). However, this short distance is not sampled robustly in nanodiscs, suggesting that this conformation is not as stable and therefore could be biased by the detergent micelles. The presence of a fully helical TM5i in the LCP crystal structure supports the notion that unwinding may either be detergent-induced or, less plausibly, reflect a higher-energy intermediate.

Consistent with the former interpretation, the partially unwound TM5i conformation was not sampled by the AF2 models. However, the cluster analysis uncovered a population of structures (Cluster 5) that adopted an elongated TM4-TM5 loop because of TM5i unwinding, but at its N-terminus as opposed to the middle of the helix as observed in 4US3 (*SI Appendix*, Fig. S10). Interestingly, hSERT cryo-EM structures show a similar effect for the IF conformation though the unwound region of TM5i extends further than observed in the MhsT models (19). This region displays variable density, strongly suggesting that the TM5i region is inherently dynamic, yet it does not rule out the possibility that TM5i samples an unwound conformation. Further biophysical studies on hSERT, particularly within nanodisc environments, are needed to assess the prevalence and functional relevance of TM5i unwinding in the context of the NSS transport cycle.

### Mechanism of ligand-dependent alternating access of MhsT

The synergistic interpretation of the DEER data in the context of SPEACH_AF2 models stimulates a transport mechanism for MhsT. Central to its alternating access is the movement of the bundle helices relative to the scaffold. On the extracellular side, coupled movements of TM1b and TM10 ensure the closure of the substrate binding site whereas intracellular movements traced by bundle helices TM1a, TM2, and TM7, along with large displacement of TM5i, facilitate the intracellular opening. Additionally, multiple TMs exhibit distinct shifts within both the bundle and the scaffold, arguing for deviation from strict rigid body transitions. Specifically, the dynamic behavior of extracellular TM8 in lipid bilayers and the relative distance changes between TM1 and TM2 on the intracellular side stand out as the most prominent examples. The SPEACH_AF2 models bin these structural changes into distinct clusters, seven were selected to sample the full range of conformational space.

The ligand dependence of the DEER distance changes demonstrates how ions and substrate stabilize conformational states of MhsT (Fig. 8). In the absence of ligands, i.e. the apo condition, an equilibrium between OF, IF, and occluded conformations is observed. While the OF state is less prevalent under apo and Na^+^ conditions, it remains accessible for ion and substrate binding. In our model, the binding of Na^+^ shifts MhsT towards a more occluded state. Binding of Trp further constricts the extracellular pathway and promotes intracellular fluctuations between occluded and IF conformations that facilitate cytoplasmic release of cargo. While our results emphasize the roles of TM1, TM5, TM6, and TM10 in the formation of distinct conformations, coordinated coupling of other helices, such as TM7 and TM8, contribute to the functional cycle. Given TM8’s crucial role in coordinating the Na2 site, our model lends support to earlier hypotheses generated from MD simulations of MhsT that suggest a TM5-TM8 interaction within the membrane could enable the release of Na2, leading to intracellular opening (26).

**Fig. 8.**
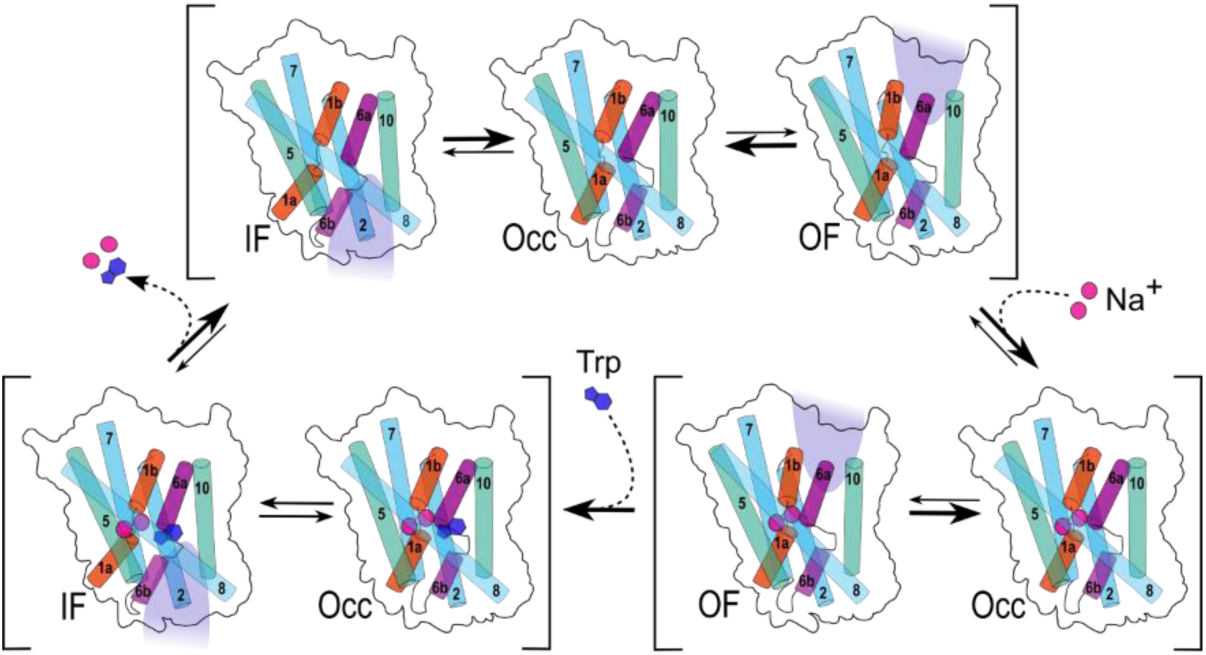
Schematic representation of the MhsT mechanism of transport. In the apo state, MhsT fluctuates between IF, OF, and occluded conformations. Binding of Na+ shifts the equilibrium towards an occluded state, prompting TM1b to move towards the extracellular scaffold while TM1a, TM5i, and TM7 shift towards the intracellular scaffold. Upon Trp binding, TM1b, TM6a, and TM10 move towards the scaffold, signaling extracellular closure, accompanied by a decrease TM8’s dynamics. Concurrently, TM5i and TM7, along with TM1a and TM2, shift away from the scaffold to support IF conformations, which are key for ion and substrate release.

### Conservation and divergence of structural motifs mediating alternating access in NSS family members

MhsT’s conformational changes have commonalities with those reported for other NSSs, with MhsT showing a closer similarity to hSERT than to LeuT. Coupled movements of key structural motifs, such as TM1b/TM6a, TM7/EL4, and TM6b/TM7, are associated with conformational changes in all three transporters (19, 31). In contrast, TM1a, TM2, TM5, and TM10 undergo movements in both MhsT and hSERT that are not observed in LeuT. Despite that TM1a is highly dynamic in LeuT, its DEER data reveal a lack of ligand-dependent changes (31), contrasting with observations in MhsT and inferences drawn from hSERT structures. While the movement of TM1a in MhsT is significantly less pronounced than in hSERT, the IF cryo-EM structure of the latter in nanodiscs lacks adequate resolution for this region. Instead, the mapping was performed from an alignment of the detergent IF structure that has high resolution of TM1a (20). Other than the ambiguous changes involving TM5i discussed above, TM5 seems to undergo similar movements in MhsT and hSERT.

TM10 exhibits more extensive movements in MhsT compared to hSERT. Given that MhsT has only 11 TMs, reduced dynamics of TM10 in hSERT may be a consequence of packing interactions with TM12 (23). On the other hand, MD simulations of hDAT have revealed that TM10 can adopt at least three distinct conformations throughout the transport cycle and function as a substrate sensor (45). Alignment of the crystal structure of OF DAT from *Drosophilia melanogaster* (dDAT) with the OF hSERT structures reveals a noticeable displacement in the positions of extracellular TMs 10 and 12 in dDAT relative to hSERT (23, 46). These observations imply that the extent of TM10’s movement may diverge between mammalian NSSs, and MhsT’s TM10 movement might more closely resemble that of DAT than SERT.

MhsT shares structural motifs associated with conformational changes not just with NSSs, but also the LeuT-fold transporter Mhp1. Mhp1, a Na^+^-dependent transporter from *Microbacterium liquefaciens*, facilitates the transport of benzyl-hydantoin through a rocking bundle mechanism (35, 47, 48). In this mechanism, the bundle and scaffold move as rigid bodies, a movement coupled to TMs 5 and 10, which serve as intracellular and extracellular gating helices, respectively. In contrast to LeuT, where the DEER data conflicted with conformational changes inferred from crystal structures (16, 31), Mhp1 conformational changes detected by DEER were by and large consistent with the model inferred from the crystal structures (35, 47, 48). While MhsT similarly shows varying degrees of coordinated movements of TMs 1, 2, 6, and 7, the bundle domain does not move as one rigid body as observed in Mhp1. MhsT similarly shows conformational changes involving intracellular TM5 and extracellular TM10, reflecting a similar gating role as observed in Mhp1. However, extracellular TM5 in MhsT also undergoes movements that are not present in Mhp1. In addition, the MhsT scaffold does not remain rigid as extracellular TM8 undergoes shifts in its dynamics associated with extracellular closure.

### MhsT’s transport mechanism compared to other LeuT-fold transporters

MhsT’s transport mechanism highlights the critical interplay between ligand binding and conformational changes that is distinct from other LeuT-fold transporters. Based on earlier studies on LeuT and Mhp1 (31, 35), the presence of Na^+^ was demonstrated to favor an OF conformation in NSSs. This hypothesis was consistent with HDX findings that suggest Na^+^ stabilizes an OF state in dDAT (49). However, recent cryo-EM structures of nanodisc-reconstituted hSERT challenge this notion by revealing Na^+^-bound structures in occluded and IF conformations instead of the anticipated OF conformation (20). Despite the absence of OF cryo-EM structures, the authors propose that hSERT still samples a conformational equilibrium when bound to Na^+^, which importantly includes OF states. MhsT similarly appears to favor occluded conformations when bound to Na^+^, especially within nanodisc environments where OF conformations are sampled less frequently, and the transporter’s dynamics permit a higher degree of fluctuation between conformations.

Na^+^ and substrate binding predominantly stabilize occluded conformations in LeuT and Mhp1 (31, 35), necessitating low-probability transitions for the release of Na^+^ and substrate from the intracellular side. In contrast, MhsT samples a mixture of occluded and IF conformations in the presence of Na^+^/Trp with lower energy barriers, likely potentiating ion and substrate release. A similar pattern of substrate-induced destabilization of intracellular structural motifs has been captured in dDAT and hSERT with HDX studies (40, 49).

However, the narrative became more nuanced as a consequence of the release of hSERT cryo-EM structures in nanodiscs, bound to Na^+^ and serotonin, which revealed both occluded and OF conformations (20). These structures challenge previous assumptions about NSS dynamics by providing static snapshots that question established models of the transport process (40, 50). This raises intriguing questions about how static structures and dynamic studies can be integrated to form a comprehensive understanding of NSS transport mechanisms.

In conclusion, our study illuminates the conformational dynamics of MhsT through a novel integration of DEER spectroscopy and AlphaFold2 conformational ensembles, revealing critical structural motifs and the modulatory role of lipid bilayers in the energetics of the transporter. These insights not only reinforce MhsT’s value as a valuable model for NSS research but also elucidate common motifs and divergent behaviors in conformational changes across the NSS family. Moreover, the integration of experimental and computational methodologies in our work establishes a comprehensive framework for evaluating the conformational dynamics of transporters, setting a precedent for future investigations into their mechanisms and evolutionary relationships.

## Materials and Methods

### Site-Directed Mutagenesis

Codon-optimized *mhsT* (Genscript) was cloned into pET19b vector with an inducible T7 promoter encoding an N-terminal 10-His tag. Single-and double-cysteine mutants were generated in the MhsT wild-type (WT) background, which is devoid of native cysteines using site-directed mutagenesis (NEB Q5) using primers designed from NEBaseChanger. MhsT mutants were confirmed by Sanger sequencing using T7 forward and reverse primers.

### Expression, Purification, and Labeling of MhsT

*Escherichia coli* C43 (DE3) cells were transformed with the pET19b vector encoding *mhsT*. Transformant colonies were used to inoculate Luria-Bertani (LB) media (LabExpress) containing 0.1 mg/mL ampicillin (Gold Biotechnology). Overnight colonies were grown at 34 °C for approximately 16 hours and used to inoculate minimal medium A at a 3:200 dilution. Cultures were incubated at 37 °C until the absorbance at 600 nm reached 0.7-0.9 at which point 1 mM isopropyl β-d-1-thiogalactopyranoside (IPTG) (Gold Biotechnology) was added, and the cultures continued incubating overnight at 18 °C. Cells were harvested by centrifugation at 5,000xg and were resuspended in 40 mL of lysis buffer (100 mM KPi, pH 7.5) supplemented with 1 mM phenylmethylsulfonyl fluoride (PMSF) (Gold Biotechnology). Cells were lysed by sonication on ice with continuous stirring. Following centrifugation at 10,000xg, the supernatant was collected, supplemented with 1 mM dithiothreitol (DTT) (Gold Biotechnology), and subsequently centrifuged at 200,000xg for 1.5 hours to isolate the membranes.

Membrane pellets were solubilized in resuspension buffer (50 mM Tris/Mes, 10% glycerol, 200 mM NaCl, 0.5 mM DTT, pH 7.5) containing 1.3% DM (Anatrace) and incubated on ice with continuous stirring for 1 hour. Insoluble material was pelleted by centrifugation at 200,000xg for 30 minutes. The resulting supernatant was incubated with 1.0 mL Ni-NTA Superflow resin (Qiagen) at 4 °C for 2 hours with rotation. The MhsT-bound resin was washed with 5 bed volumes of resuspension buffer supplemented with 50 mM imidazole and 0.15% DM. MhsT was eluted with resuspension buffer supplemented with 300 mM imidazole and 0.15% DM.

Following affinity purification, MhsT mutants undergoing spin-labeling were supplemented with 60 mM Mes to decrease the pH of the sample to 7.0. purified MhsT mutants were labeled with 10-fold molar excess of (1-Oxyl-2,2,5,5-tetramethyl-3-pyrroline-3-methyl)methanethiosulfonate (MTSSL) (Enzo Life Sciences) per cysteine followed by two rounds of 5-fold molar excess MTSSL at room temperature in the dark for a period of 3 hours. Following labeling, the protein sample was placed on ice at 4 °C overnight. MhsT was concentrated to 1-2 mL using 50,000 Dalton MWCO filter concentrators (Millipore) and centrifuged at 105,000xg for 20 minutes. Excess spin label was removed by size-exclusion chromatography (SEC) over a Superdex200 Increase 10/300 GL column (Cytiva) into detergent SEC buffer (50 mM Tris/Mes, 0.15% DM, 10% glycerol, pH 7.2). Peak fractions of purified MhsT typically eluted between 12.5-14 mL, and the concentration of MhsT was quantified by absorbance at 280 nm (ε= 98,760 M^-1^ cm^-1^). Purified MhsT was then allocated for the following experiments: proteoliposome reconstitution for transport assays, DM DEER sample preparation, or nanodisc reconstitution for DEER sample preparation.

### Reconstitution of MhsT into Proteoliposomes

*E. coli* polar lipids (Avanti Polar Lipids) were dissolved in chloroform and evaporated by N_2_ gas while stirring. Lipids were resuspended to 20 mg/mL in 100 mM KPi, pH 6.5 supplemented with 1.5% β-OG (Anatrace) and then dialyzed using a 3,500 Dalton MWCO dialysis cassette (ThermoFisher) into 100 mM KPi, pH 6.5 overnight at 4 °C to remove OG (21). Preformed liposomes were frozen in liquid N_2_ and stored in aliquots at-80°C.

Prior to reconstitution, preformed liposomes were thawed, diluted to 5 mg/mL lipids with inside buffer, and extruded using an Avestin LiposoFast extruder. Preformed lipids were extruded through a 100 nm membrane filter (Avestin) inserted between two polyester 10 mm drain discs (Whatman). Following extrusion, 0.11% Triton-X-100 was added to the preformed liposomes, and the mixture was vortexed. Purified MhsT (or buffer for control liposomes) was added to the lipid-detergent mixture at 1:150 (w/w), and the reaction was allowed to incubate with gentle agitation at room temperature. Following an incubation period of 15 minutes, a total of 240 mg/mL Bio-Beads SM-2 (BioRad) were added incrementally to the reaction over the next two hours to remove detergent. The reaction continued to incubate with gentle agitation overnight at 4 °C. Following the removal of Bio-Beads, proteoliposomes (or control liposomes) were ultracentrifuged at 320,000xg for 45 minutes and subsequently resuspended to a final lipid concentration of ∼5 mg/mL with inside buffer. MhsT protein concentration was quantified by densitometry (ImageJ) of SDS/PAGE gels stained with Oriole fluorescent dye (Biorad) using a standard curve of detergent-purified MhsT as a reference. Samples were aliquoted and frozen in liquid N_2_ prior to conducting transport assays.

### Transport Assays

In preparation of transport assays, proteoliposome-reconstituted MhsT samples were thawed at room temperature and extruded in the same manner as described above for the lipids. The transport activity of proteoliposome-reconstituted MhsT was measured by the accumulation of [^3^H]-Trp (18 Ci/mmol; American Radiolabeled Chemical, Inc). For both experiments, reactions were initiated by the addition of proteoliposomes containing 30 ng MhsT (or the equivalent volume for control liposomes) to 100 µL assay buffer (10 mM Tris/Mes, 150 mM NaCl, pH 8.5). Time-dependent uptake was measured at a [^3^H]-Trp concentration of 0.1 µM, whereas the concentration-dependent accumulation of [^3^H]-Trp (ranging from 0.1 µM to 20 µM) was measured for periods of 10 seconds. Reactions were quenched by adding cold stop buffer (100 mM KPi, 100 mM LiCl, pH 6.0) and vacuum filtered through 0.45 µM mixed cellulose ester membrane filters (Millipore). Filters were subjected to scintillation counting (Hides SL300). MhsT-specific uptake activity was determined by subtracting the accumulation of [^3^H]-Trp measured in control liposomes lacking MhsT from the accumulated [^3^H]-Trp measured in MhsT-containing proteoliposomes under the same experimental conditions. For the concentration-dependent [^3^H]-Trp uptake measurements, the 10-s uptake data were corrected for the data determined at the zero timepoint. Decays per minute (dpm) were transformed to pmol using known amounts of [^3^H]-Trp. Data were analyzed using Michaelis-Menten fitting in GraphPad Prism 10. Data are shown as mean ± S.E.M. (n≥3), and kinetic constants are the mean ± S.E.M. (n≥3) of the fit. Due to the large sample size, MhsT mutants were assayed in nine batches, and MhsT-WT-containing proteoliposomes served as reference for each batch. The concentration of Trp present in the Na^+^/Trp DEER sample exceeds the K_m_ for each mutant by at least tenfold (*SI Appendix*, Table S1). With the exception of the 251-424 mutant, all mutants retain a V_max_ ≥ 30% of WT. The lower V_max_ for 251-424 is likely due to aggregation of the mutant as indicated by the broad, long-distance component in the DEER data.

### Preparation of DM DEER Samples

Following SEC, MhsT fractions were concentrated using 50,000 Dalton MWCO filter concentrators (Millipore) and supplemented with glycerol to bring the final glycerol concentration to 23% for cryoprotective purposes. The subsequent concentration of MhsT was quantified by absorbance at 280 nm (ε = 98,760 M^-1^ cm^-1^). The spin label concentration was evaluated by continuous-wave electron paramagnetic resonance (CW-EPR). The spin labeling efficiency of the sample was determined by assessing the ratio of the experimental spin label concentration to the theoretical maximum spin label concentration. DEER samples were prepared under three biochemical conditions: apo, Na^+^, and Na^+^/Trp. Each sample was diluted 20% using a combination of detergent SEC buffer, NaCl, and Trp as required for each condition. Specifically, 200 mM NaCl and 1 mM Trp were added to samples as necessary. The DEER samples were frozen in liquid N_2_ prior to DEER spectroscopy.

### Reconstitution of MhsT into Nanodiscs & DEER Sample Preparation

A 3:1 (w/w) mixture of *E. coli* polar lipids and egg L-α-phosphatidylcholine (Avanti Polar Lipids) was dissolved in chloroform, evaporated on a rotary evaporator, and desiccated under vacuum overnight in the dark. Following resuspension of the lipid mixture to a concentration of 20 mM with lipid resuspension buffer (200 mM Tris/Mes, pH 7.2), the lipids were homogenized by 10 freeze-thaw cycles and stored in small aliquots at-80°C. MSP1D1E3 was purified as previously described (51). Purified MSP1D1E3 was concentrated using 10,000 Dalton MWCO filter concentrators (Millipore) towards an optimal concentration of 0.6-0.8 mM as determined by the absorbance at 280 nm (ε = 29,910 M^−1^ cm^−1^). MSP was diluted with 5% glycerol and stored in aliquots at-80°C.

Reconstitution of MhsT into nanodiscs proceeded with the addition of spin labeled MhsT mutants in DM micelles to a mixture of homogenized lipids, MSP1D1E3, and sodium cholate. A molar ratio of 1:8 MhsT:MSP1D1E3, 1:50 MSPD1E3:lipid, and 1:4 lipid:cholate was used for the reaction. The reconstitution reaction was allowed to incubate with gentle agitation at 4°C. Following an incubation period of 1 hour, a total of 400 mg/mL Bio-Beads SM-2 (BioRad) were added incrementally over the next two hours to remove detergent. The reaction continued to incubate with gentle agitation overnight at 4°C. The next day, 200 mg/mL Bio-Beads were added to the reaction that was allowed to incubate for an additional hour. Bio-Beads were removed by filtering the sample using a 0.20-µm filter (Corning). MhsT-reconstituted nanodiscs were then separated from empty nanodiscs by SEC into nanodisc SEC buffer (200 mM Tris/Mes, 10% glycerol, pH 7.2). The relative protein:MSP ratio in the obtained fractions was quantified by ImageJ of SDS/PAGE gels. Suitable fractions were concentrated using 100,000 Dalton MWCO concentrators (Millipore) and supplemented with glycerol to bring the final glycerol concentration to 23% for cryoprotective purposes. The MhsT protein concentration and associated reconstitution efficiency was determined from the spin label concentration characterized by CW-EPR and the spin labeling efficiency as calculated above. DEER samples were prepared under three biochemical conditions: apo, Na^+^, and Na^+^/Trp. Each sample was diluted 20% using a combination of nanodisc SEC buffer, NaCl, and Trp as required for each condition. Specifically, 200 mM NaCl and 1 mM Trp were added to samples as necessary. The DEER samples were frozen in liquid N_2_ prior to DEER spectroscopy.

### CW-EPR and DEER Spectroscopy and Data Analysis

CW-EPR of spin-labeled MhsT samples were collected at room temperature using a Bruker EMX spectrometer operating at X-band frequency (9.5 GHz), 10-mW incident power, and a 1.6 G modulation amplitude. DEER spectroscopy of doubly labeled MhsT samples were collected at 50 K using an Elexys E580 EPR spectrometer operating at Q-band frequency (33.9 GHz) with a dead-time-free four-pulse protocol. The observe pulse lengths were 6 or 8 ns (π/2) and 12 or 16 ns (π), and the pump pulse length was 40 ns. The frequency separation between the observe and pump pulses was set as 79.24 MHz.

Raw DEER decays were analyzed using home-written software operating in a Matlab environment as previously described (52, 53). In this program, the distance distributions are assumed to consist of a sum of Gaussians. While previous studies have used global analysis to evaluate how different biochemical conditions shift the population of the distances within the distribution, the individual fits of the DEER decays were used in our evaluation of MhsT to reduce biases associated with the assumption that the center and width of the Gaussians were shared across conditions.

### SPEACH_AF Model Generation

We utilized SPEACH_AF, an in-silico mutagenesis method that leverages AlphaFold2 (AF2) to simulate protein ensembles. Alanine substitutions were introduced across the multiple sequence alignment (MSA) using a 5 or 10 amino acid sliding window approach. For each set of mutations, 3 models were generated each from the five distinct AF2 modelers, resulting in a total of 1,920 models. These models were then screened using average predicted local distance difference test (pLDDT) scores and principal component analysis (PCA), with all MhsT models successfully passing these filters. PCA was conducted using ProDy software (54–56).

To identify collective variables, we analyzed the root-mean-square fluctuation (RMSF) of all models with ProDy, pinpointing a pair of high and low RMSF values indicative of MhsT’s extracellular or intracellular sides opening and closing. The RMSF analysis resulted in selecting residue pairs 41-121 for the extracellular side and 16-343 for the intracellular side. We applied K-means clustering (via the sklearn library) to these collective variables, increasing the number of clusters until models with the most extreme values for the 16-343 variable were in their own cluster, which occurred at seven clusters.

To find a representative model for each cluster, we computed the root-mean-square deviation (RMSD) of each model against others within the same cluster using ProDy. The model with the lowest average RMSD was designated as the representative for its respective cluster. Finally, per-residue RMSD comparisons between the representative models of each cluster were performed in PyMOL. This involved aligning each model pair and then calculating the Cα-Cα distances between corresponding residues.

## Acknowledgments

We thank Dr. Derek P. Claxton for critical review of this manuscript. We also thank Mr. Michael Mohan for his assistance in creating mutants and protein expression and purification. This work was supported by the National Institutes of Health, through National Institute of Neurological Disorders and Stroke F31 (F31NS125911) and National Institute of General Medical Sciences for the Molecular Biophysics Training Grant T32 (T32GM008320).

## Author Contributions

A.C.S. and H.S.M. designed research; A.C.S., E.G., and R.A.S. performed research; A.C.S., E.G., M.Q., and R.A.S. analyzed data; and A.C.S., R.A.S., and H.S.M. wrote the paper.

## Classification

Biological Sciences, Biophysics and Computational Biology

## Supplemental Figures and Tables

**Fig. S1.**
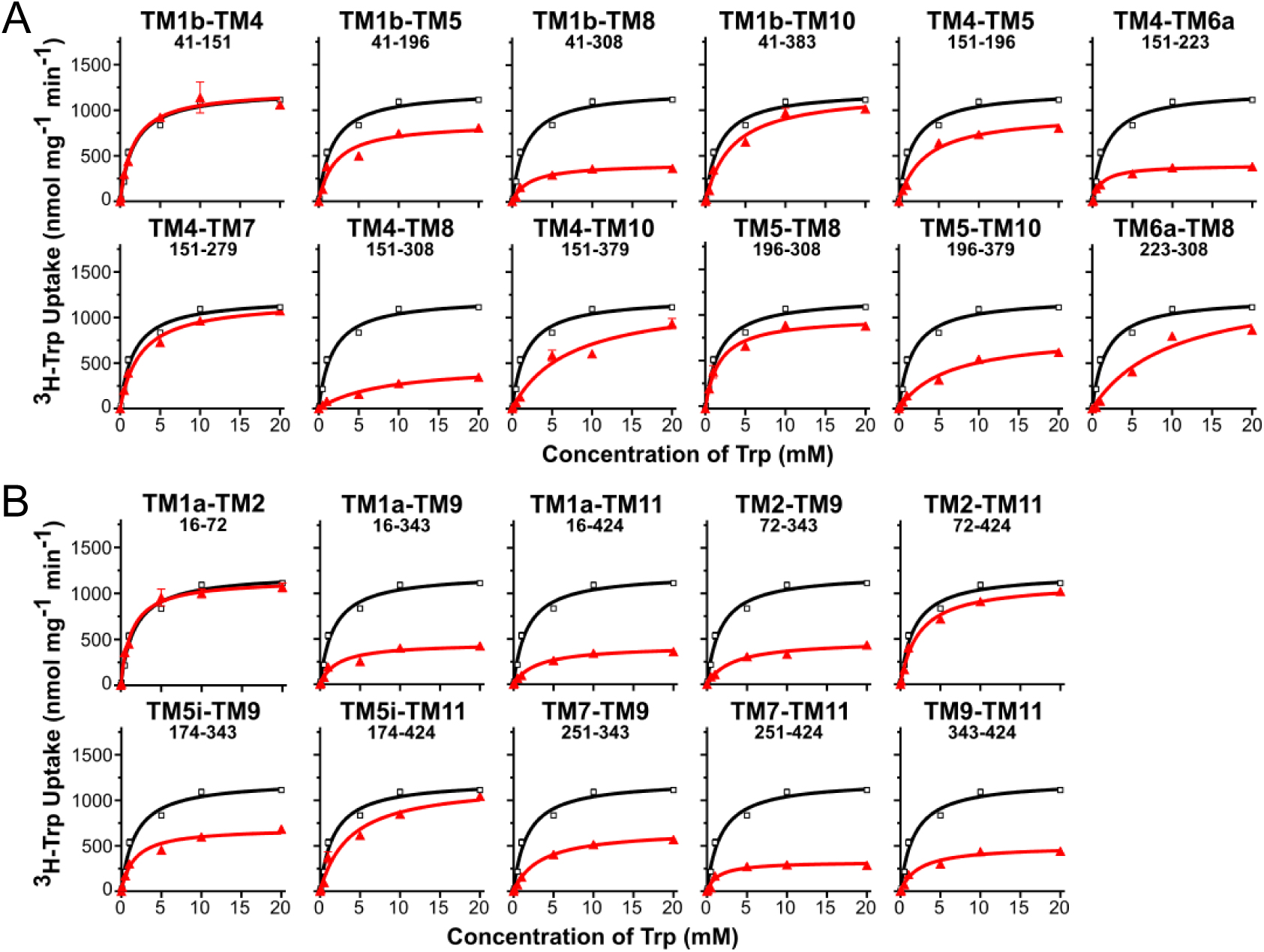
Uptake of ^3^H-Trp for spin-labeled MhsT double mutants in proteoliposomes. Uptake curves were fitted to the Michaelis-Menten equation and plotted for extracellular (*A*) and intracellular (*B*) mutants are shown in red, WT in black for comparison, and error bars indicate standard error of the mean (*n*=3).

**Table S1.** Kinetic parameters for spin-labeled MhsT double mutants.

Extracellular cysteine pairs.
| Mutant | $V_{\max} \pm \text{S.E.M.}$<br>(nmol mg<sup>-1</sup> min<sup>-1</sup>) | | | $K_M \pm \text{S.E.M.}$<br>($\mu\text{M}$) | | |
| --- | --- | --- | --- | --- | --- | --- |
| MhsT WT | 1217 | ± | 65 | 1.71 | ± | 0.37 |
| 41-151 | 1152 | ± | 41 | 1.34 | ± | 0.20 |
| 41-196 | 862 | ± | 82 | 2.09 | ± | 0.76 |
| 41-308 | 413 | ± | 22 | 2.13 | ± | 0.44 |
| 41-383 | 1205 | ± | 87 | 3.33 | ± | 0.81 |
| 151-196 | 966 | ± | 58 | 3.26 | ± | 0.67 |
| 151-223 | 399 | ± | 15 | 1.15 | ± | 0.18 |
| 151-279 | 1186 | ± | 53 | 2.50 | ± | 0.41 |
| 151-308 | 489 | ± | 62 | 8.33 | ± | 2.47 |
| 151-379 | 1213 | ± | 179 | 7.14 | ± | 2.63 |
| 196-308 | 1003 | ± | 50 | 1.74 | ± | 0.34 |
| 196-379 | 827 | ± | 82 | 6.29 | ± | 1.64 |
| 223-308 | 1371 | ± | 256 | 10.03 | ± | 4.09 |

**B** Intracellular cysteine pairs.
| Mutant | $V_{\max} \pm \text{S.E.M.}$<br>(nmol mg<sup>-1</sup> min<sup>-1</sup>) | | | $K_M \pm \text{S.E.M.}$<br>($\mu\text{M}$) | | |
| --- | --- | --- | --- | --- | --- | --- |
| MhsT WT | 1217 | ± | 65 | 1.71 | ± | 0.37 |
| 16-343 | 453 | ± | 44 | 2.14 | ± | 0.80 |
| 16-424 | 425 | ± | 13 | 2.96 | ± | 0.32 |
| 16-72 | 1224 | ± | 54 | 1.64 | ± | 0.29 |
| 72-343 | 480 | ± | 30 | 3.17 | ± | 0.69 |
| 72-424 | 1119 | ± | 49 | 2.31 | ± | 0.38 |
| 174-343 | 694 | ± | 40 | 1.61 | ± | 0.38 |
| 174-424 | 1191 | ± | 113 | 3.74 | ± | 1.14 |
| 251-343 | 673 | ± | 14 | 3.34 | ± | 0.24 |
| 251-424 | 326 | ± | 26 | 1.37 | ± | 0.46 |
| 343-424 | 498 | ± | 36 | 2.32 | ± | 0.62 |

**Fig. S2.**
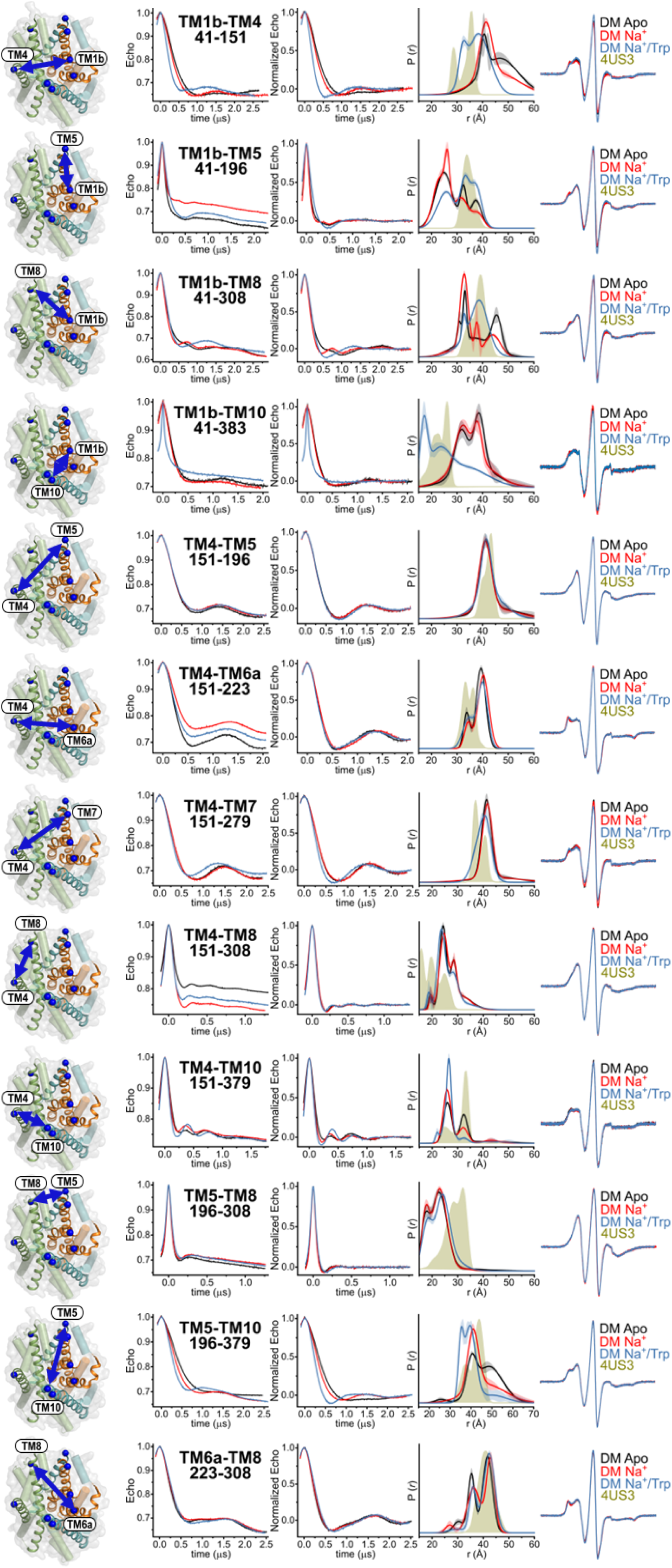
DEER analysis of extracellular spin label mutants in DM micelles under three biochemical conditions: apo (black), Na^+^ (red), and Na^+^/Trp (blue). The raw and normalized DEER data are shown alongside the fitted distributions with confidence bands as well as the CW-EPR traces as controls. The predicted 4US3 crystal structure distribution is shown for comparison.

**Fig. S3.**
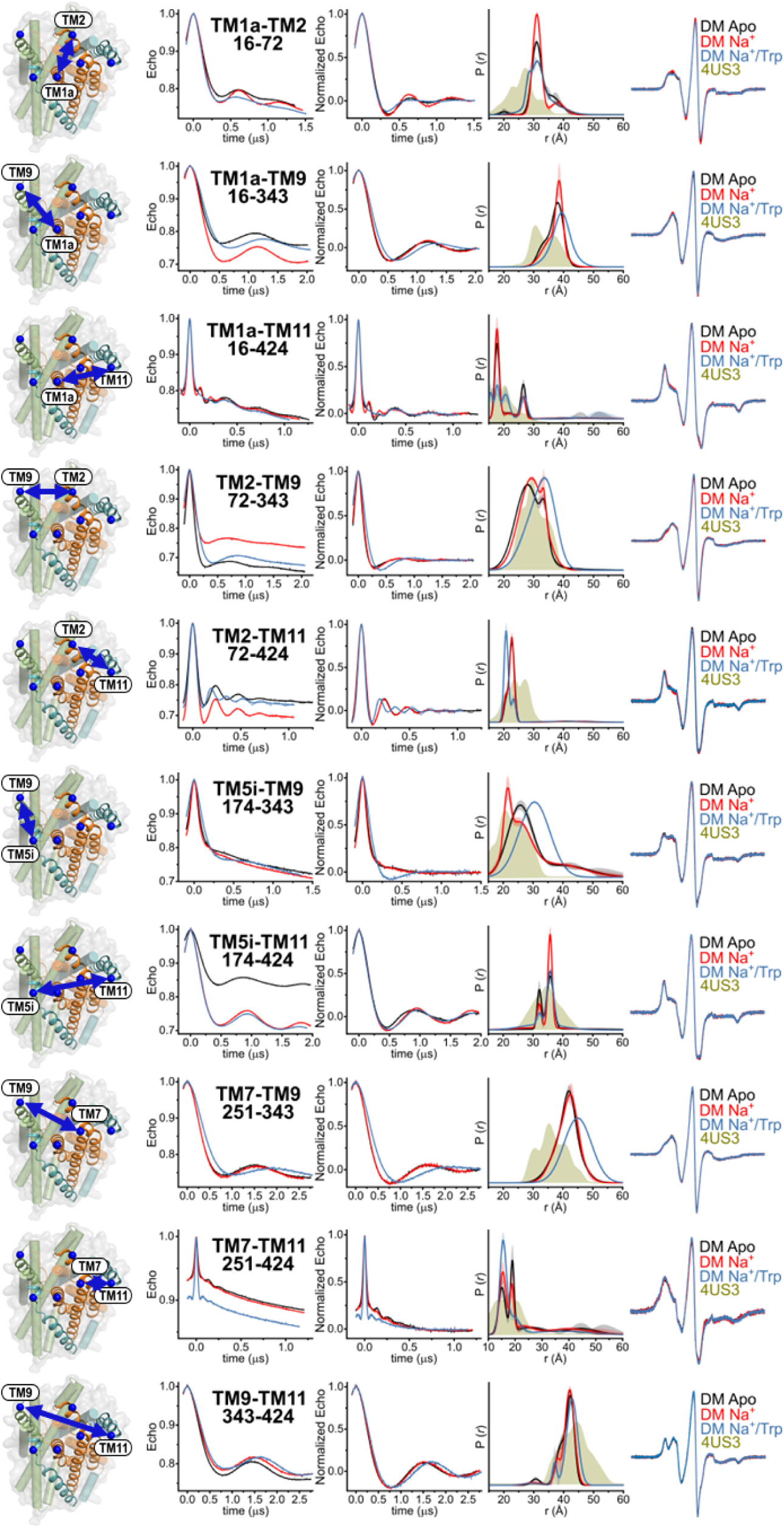
DEER analysis of intracellular spin label mutants in DM micelles under three biochemical conditions: apo (black), Na^+^ (red), and Na^+^/Trp (blue). The raw and normalized DEER data are shown alongside the fitted distributions with confidence bands as well as the CW-EPR traces as controls. The predicted 4US3 crystal structure distribution is shown for comparison.

**Fig. S4.**
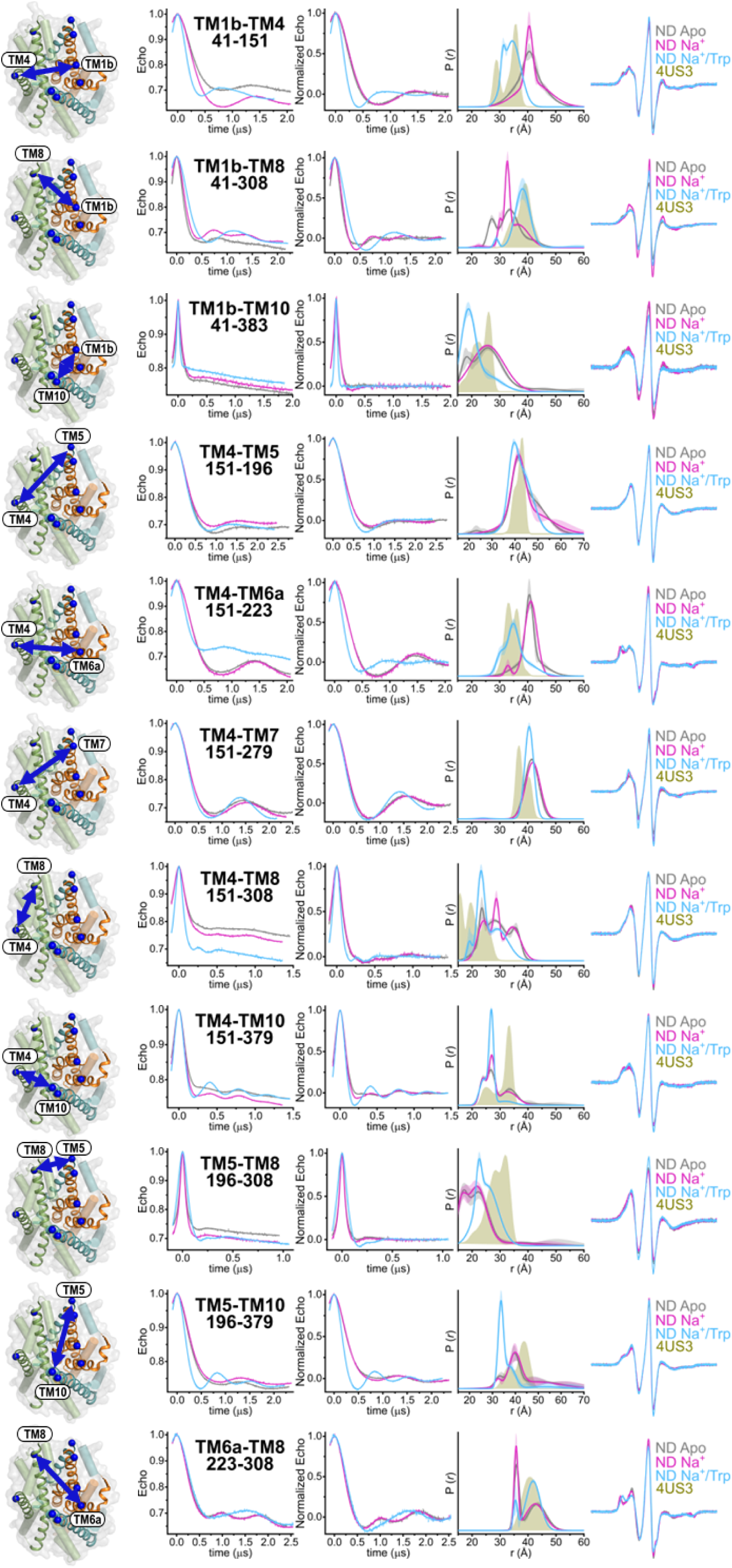
DEER analysis of extracellular spin label mutants in lipid nanodiscs (ND) under three biochemical conditions: apo (gray), Na^+^ (pink), and Na^+^/Trp (light blue). The raw and normalized DEER data are shown alongside the fitted distributions with confidence bands as well as the CW-EPR traces as controls. The predicted 4US3 crystal structure distribution is shown for comparison.

**Fig. S5.**
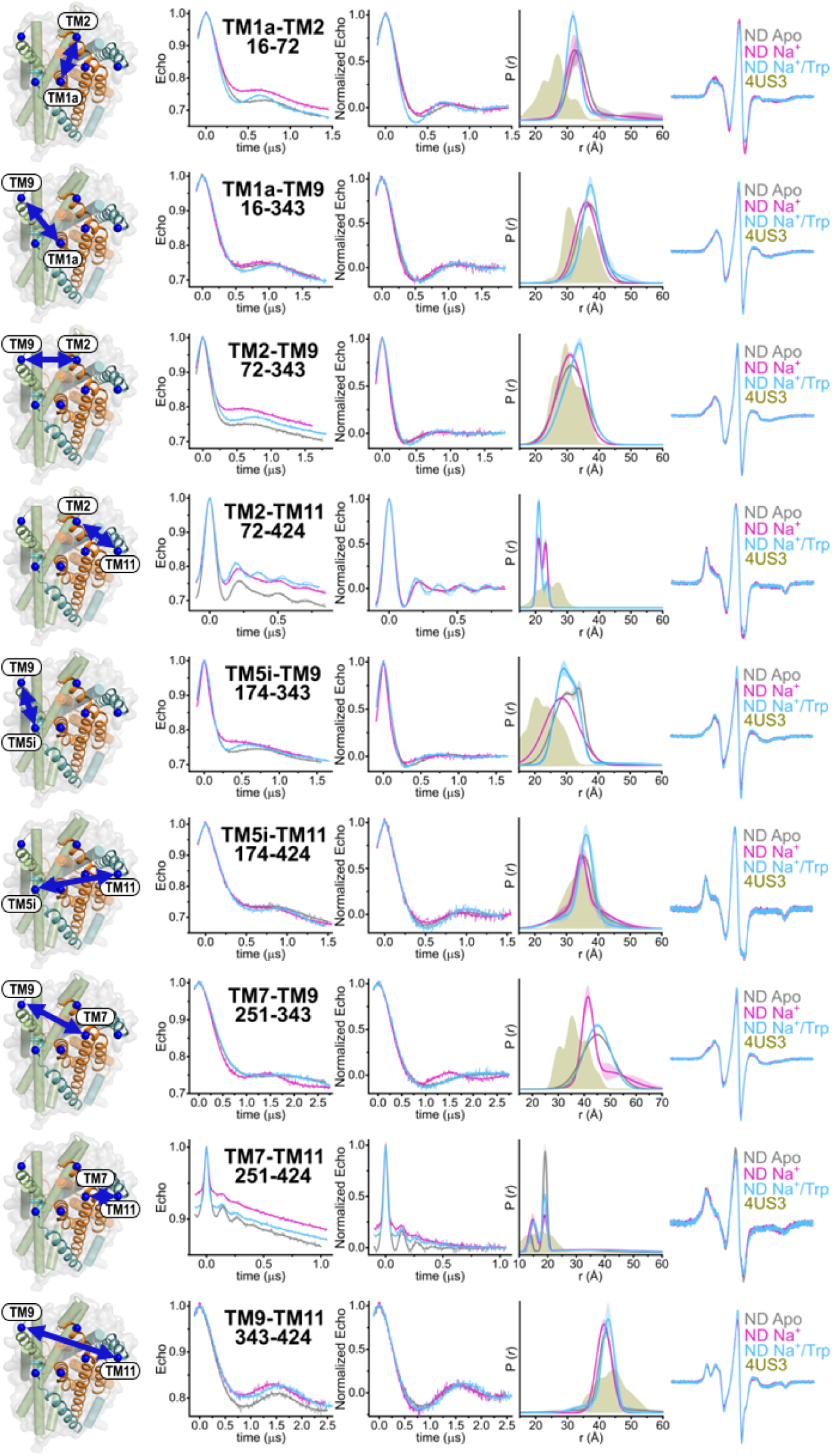
DEER analysis of intracellular spin label mutants in lipid nanodiscs (ND) under three biochemical conditions: apo (gray), Na^+^ (pink), and Na^+^/Trp (light blue). The raw and normalized DEER data are shown alongside the fitted distributions with confidence bands as well as the CW-EPR traces as controls. The predicted 4US3 crystal structure distribution is shown for comparison.

**Fig. S6.**
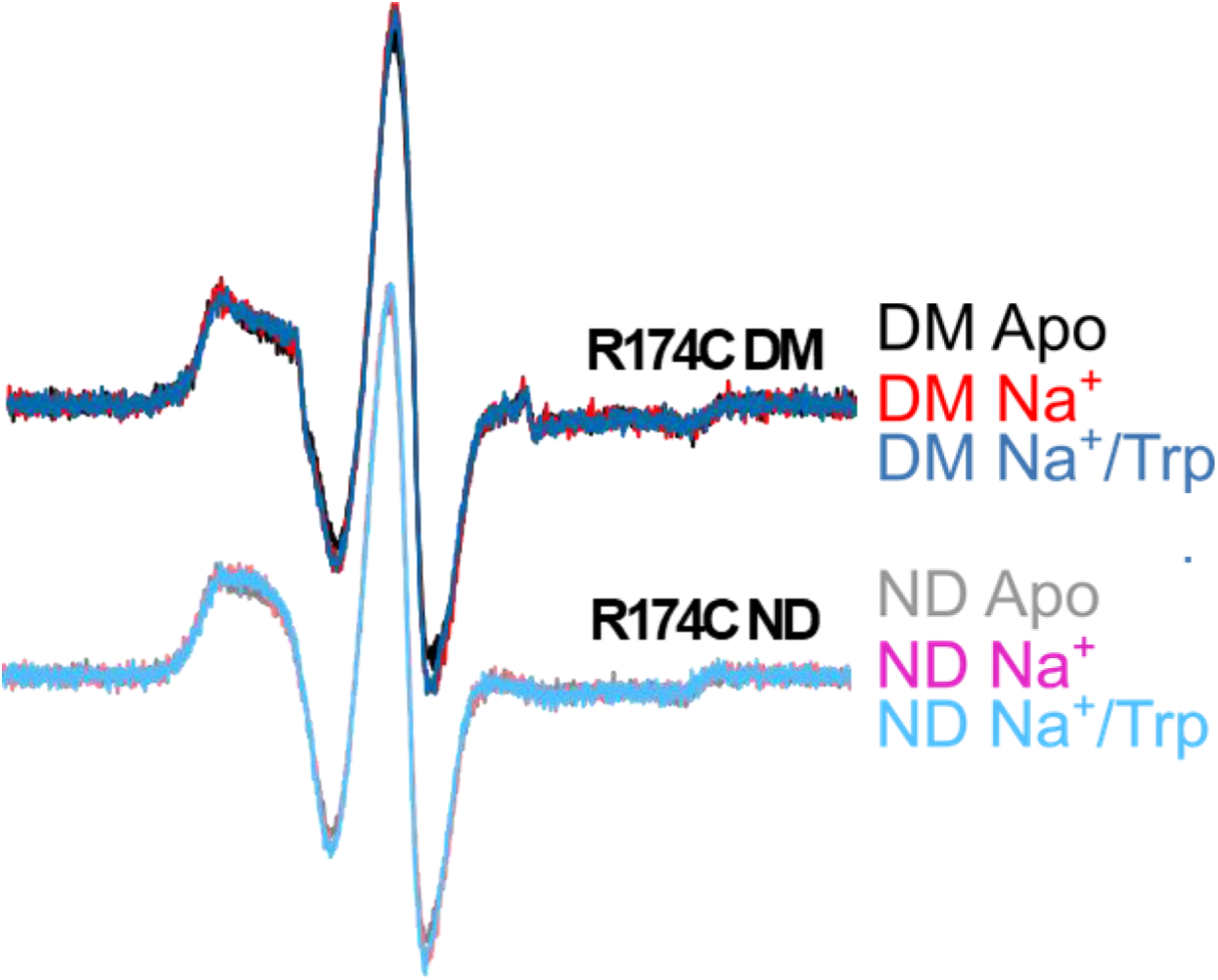
CW-EPR of singly labeled TM5i site R174C in different environments. Spectra demonstrate that the spin label is located on an ordered region of TM5i under apo, Na^+^, and Na^+^/Trp conditions in both detergent micelles (DM) and lipid nanodiscs (ND).

**Fig. S7.**
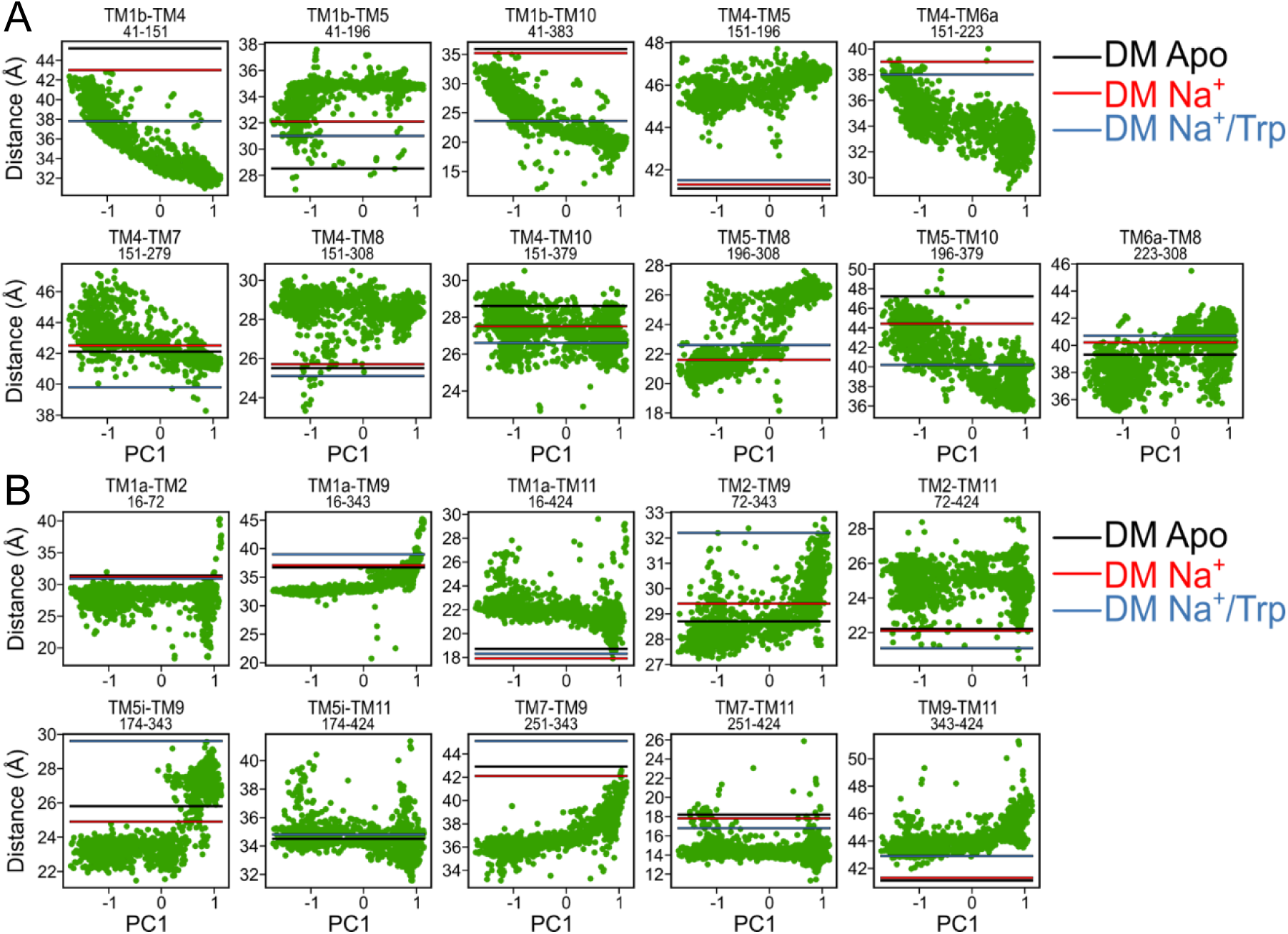
Simulated versus experimental distances along PC1 for extracellular (*A*) and intracellular (*B*) spin label pairs. Analysis of the extracellular side shows a broad range of conformational states across all of PC1 space. In contrast, conformational changes on the intracellular side are confined to the PC1 range of 0.5-1, suggesting an asymmetry in the degree of conformational changes between the two sides of the transporter under the tested biochemical conditions.

**Fig. S8.**
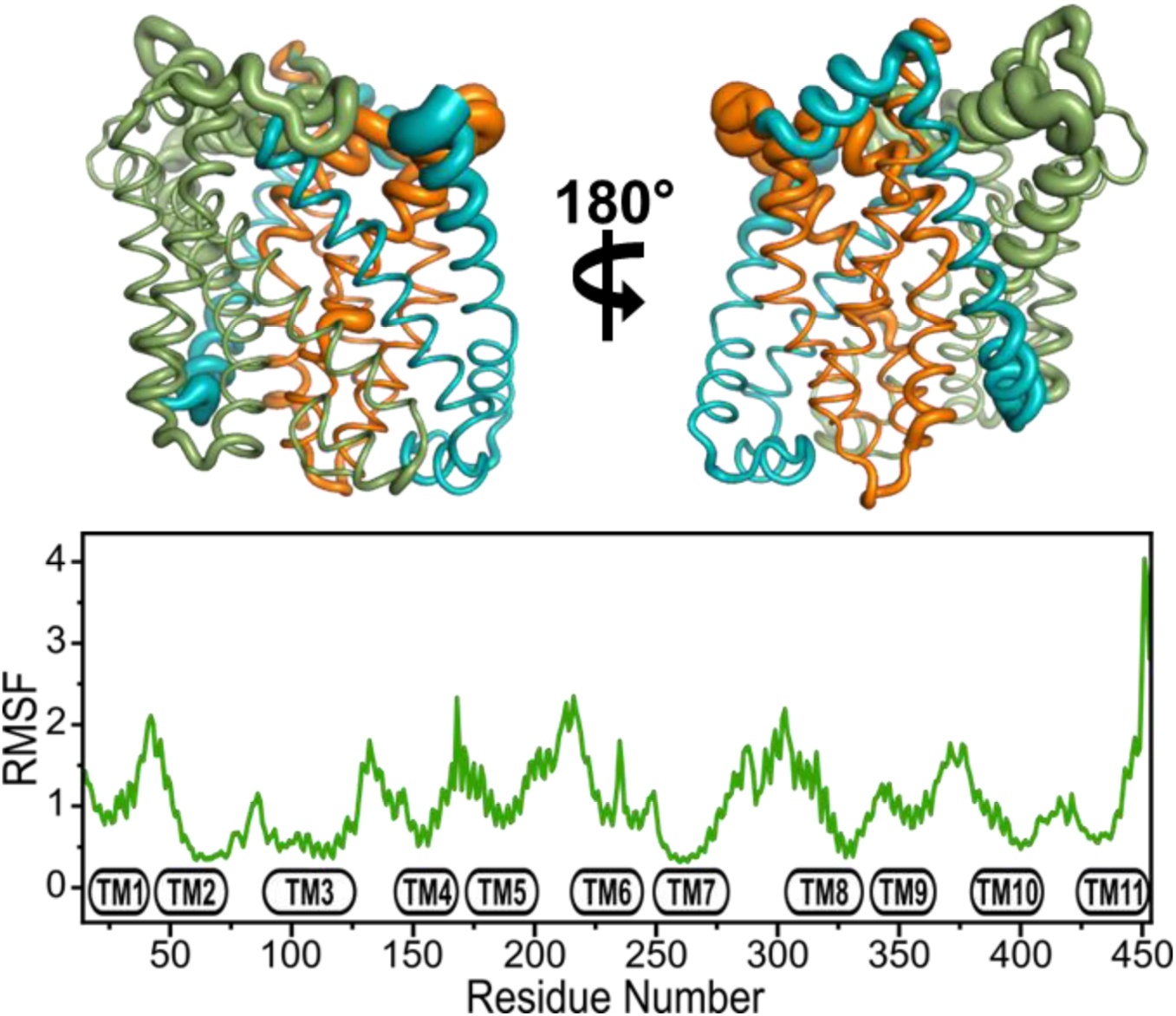
RMSF values calculated across all generated models mapped onto occluded cluster 4 model (*Top*) and plotted on a per-residue basis (*Bottom*). Cluster 4 model represents the closest structure to a doubly occluded conformation though biased towards the intracellular side and most resembles the MhsT crystal structure conformation. The scaffold domain (TM3, TM4, TM8, TM9) is shown in green, the bundle domain (TM1, TM2, TM6, TM7) is shown in orange, and TMs 5, 10 and 11 are shown in teal.

**Fig. S9.**
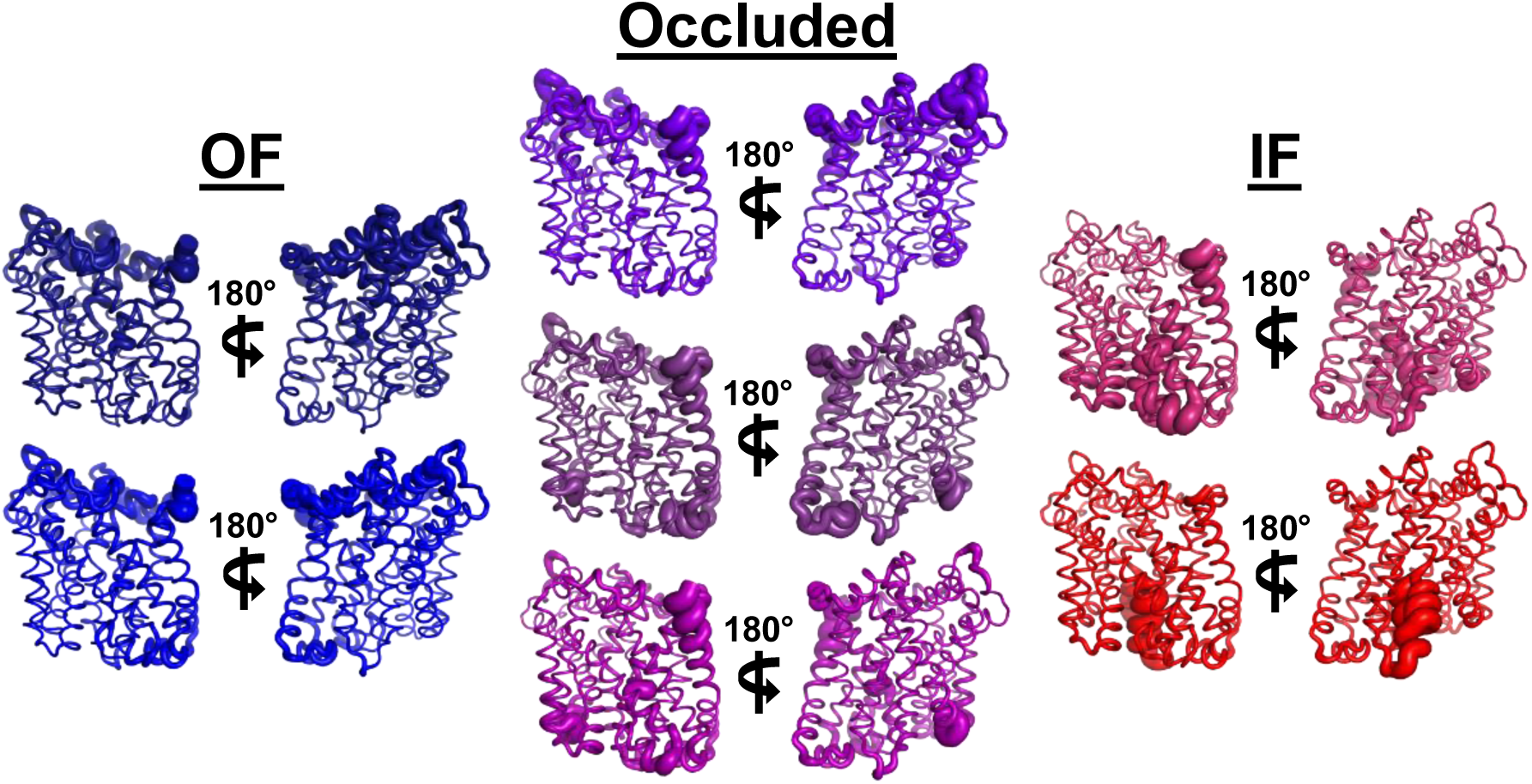
Residue fluctuation profiles for each cluster mapped onto structural models for Clusters 1-7. Clusters range from most outward-open (blue) to most inward-open (red). OF Clusters 1-2 (*Left*) and occluded OF Cluster 3 (*Middle, Top*) show higher variability compared to occluded IF Clusters 4 and 5 (*Middle*). IF Clusters 6 and 7 (*Right*) vary in their levels of their disorder: partially IF Cluster 6 exhibits larger disorder relative to the fully IF Cluster 7, with TM1 and TM6 as notable exceptions.

**Fig. S10.**
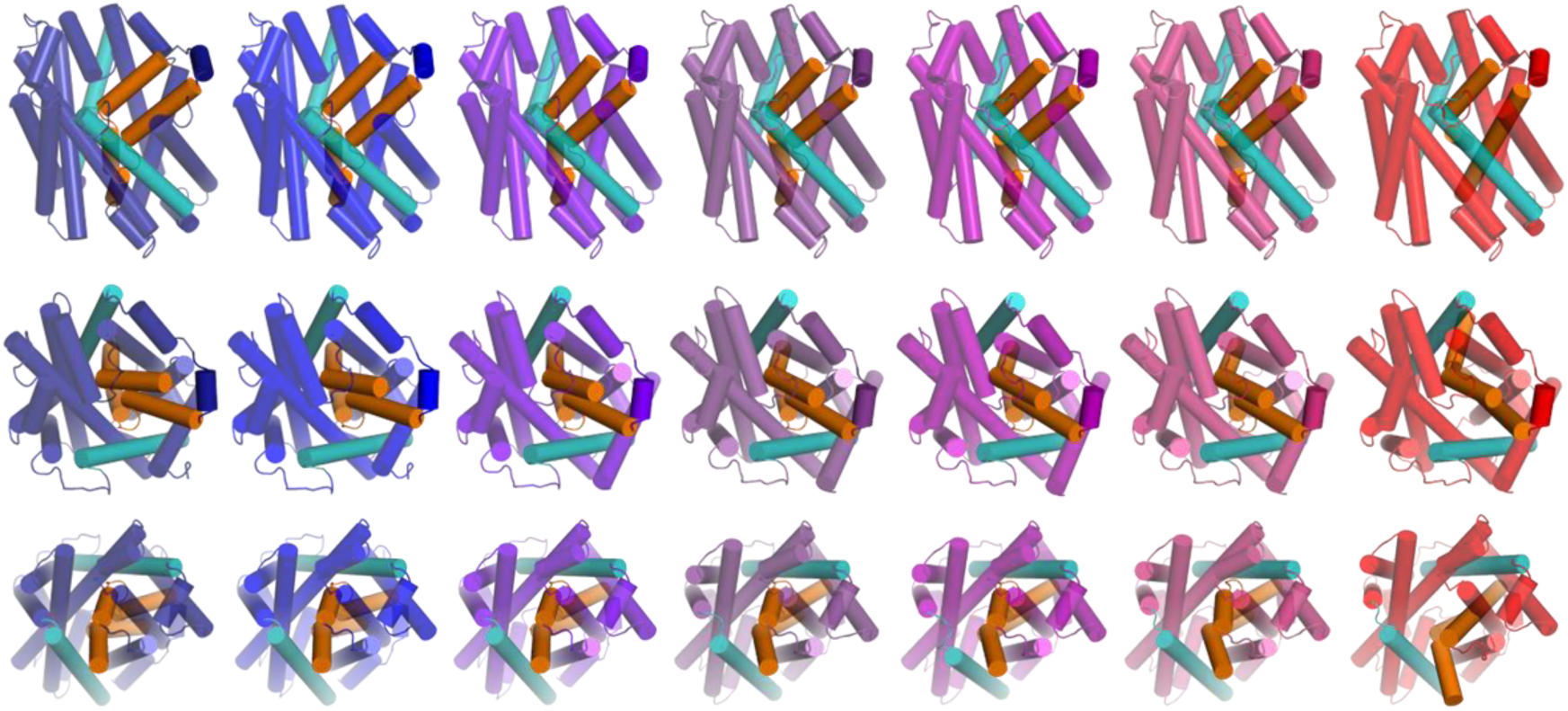
Multiple views of representative models of each cluster. Clusters range from most outward-open (blue) to most inward-open (red). Bundle TMs 1 & 6 are shown in orange while gating TMs 5 & 10 are shown in teal. Panels depicts the view from the side (*Top*), from the extracellular side (*Middle*), and from the intracellular side (*Bottom*).

**Fig. S11.**
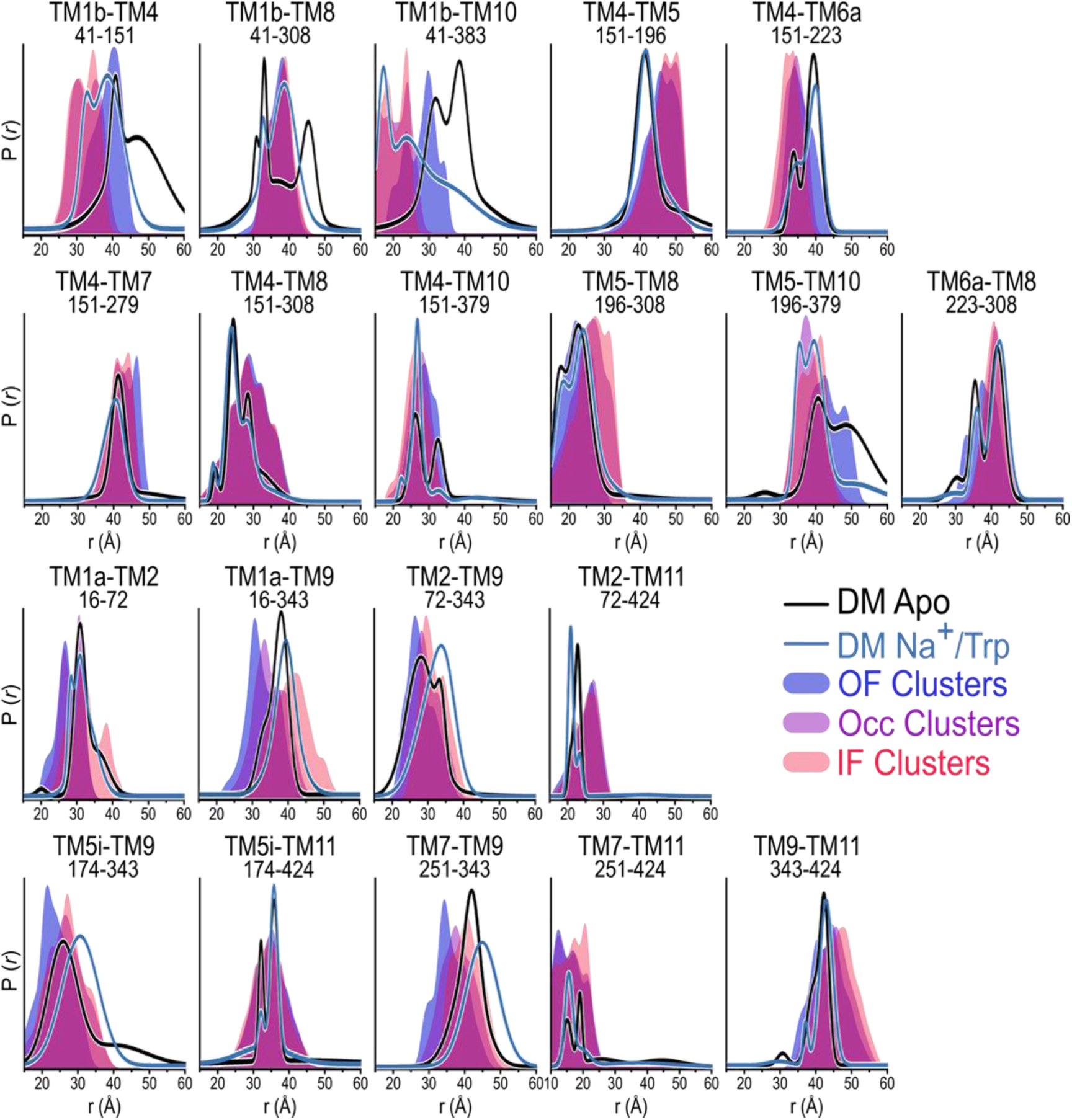
Comparison of predicted MhsT conformational state distance distributions with DEER data. Solid colors depict the averaged distance distributions for clusters classified as OF (blue), occluded (purple), and IF (red) states. Overlaid lines represent experimental DEER data for DM Apo (black) and DM Na^+^/Trp (blue) conditions, facilitating direct visual comparison between simulated spin label pairs across conformations and experimentally derived data.

